# A Single-Cell Signaling Atlas of Spinal Cord BDNF Responses Reveals Determinants Beyond Receptor Expression

**DOI:** 10.64898/2026.03.24.713745

**Authors:** Jonathon M. Sewell, Autumn C. Bissett, Grace Lee, Eli R. Zunder, Bettina Winckler, Chris D. Deppmann

## Abstract

Although the influence of Brain-Derived Neurotrophic Factor (BDNF) has been characterized across numerous neural settings, how individual cells decode this pleiotropic message into context-dependent signaling responses remains unresolved. Using highly multiplexed single-cell mass cytometry to simultaneously measure levels of 19 signaling markers and 18 cell ID markers, we constructed a temporal atlas of BDNF-induced signaling relative to two control conditions across diverse spinal cord lineages and maturation states. We demonstrate that not all cells contribute to the global BDNF response with ∼47-75% of cells having increased ERK phosphorylation at peak activation. Our analysis of 20 uniquely identified cell identities reveals that TrkB/p75NTR receptor stoichiometry sets the potential for response, but ultimately the sustained reduction of surface TrkB predicts BDNF sensitivity. Surprisingly, identical receptor profiles in distinct cell types yield fundamentally different signaling responses, indicating that cell identity acts as the final arbiter of the BDNF message. These findings reframe BDNF sensitivity as a form of prepared competence. This work thus provides a framework for understanding how intracellular context dictates the functional interpretation of neurotrophic cues.

## INTRODUCTION

Although Brain-Derived Neurotrophic Factor (BDNF) is a single molecular message, a mature homodimeric ligand (*1*), it orchestrates many different neurobiological processes, from neurodevelopmental patterning to synaptic plasticity underlying learning and memory (*2, 3*). While the canonical model suggests that the receptor expression profile dictates the response (*4, 5*), the broad range of functional outcomes across the nervous system that are attributed to a single ligand implies that the ligand-receptor interaction alone is insufficient to explain the broad functional interpretations of the BDNF message.

The interplay between the receptors TrkB and p75NTR is widely viewed as a regulatory switch that tunes the BDNF signaling response. Upon ligand binding, TrkB autophosphorylates to initiate core Ras/MAPK, PI3K/Akt, and PLC-gamma cascades (*6*), yet these pathways are qualitatively and quantitatively modified by p75NTR co-expression. This interaction can synergistically enhance TrkB affinity (*7–10*), prolong signaling duration by reducing receptor ubiquitination (*11*), or independently activate NF-kappa-B and JNK/c-Jun stress-response pathways (*2*). However, because these mechanistic ‘rules’ were largely defined in transfected cell lines or restricted primary lineages, it remains unclear whether receptor stoichiometry alone can predict signaling outcomes across the heterogeneous cell populations of the nervous system. While single-cell signaling heterogeneity is a recognized feature of other receptor tyrosine kinase (RTK) systems (*12*), a systematic mapping of how cell identity and maturation state filter the BDNF message is currently unexplored.

This knowledge gap is primarily due to the reliance on biochemical techniques, such as immunoblotting, which measure an aggregate output that is blind to cellular heterogeneity. While these bulk readouts have provided a thorough profile of the aggregate BDNF signaling cascade, they cannot uncover how a single ligand might be differentially interpreted across non-uniform subpopulations (*13*). Newer single-cell technologies offer higher resolution but present contrasting trade-offs: single-cell RNA sequencing effectively maps cell identity but fails to capture the rapid, post-translational phosphorylation events that define the immediate BDNF response (*4*). Similarly, while standard fluorescence-based flow cytometry provides single-cell signaling data, it is constrained by spectral overlap, preventing the simultaneous measurement of the broad repertoire of phosphoproteins and receptor profiles required to decode a complex signaling landscape (*14*).

To resolve this decoding problem, we employed highly multiplexed single-cell mass cytometry to construct a high-resolution time course of BDNF signaling across neural lineages and maturation states. By simultaneously measuring receptor composition, receptor surface abundance, and a broad panel of downstream phosphoproteins, our data show a highly filtered signaling landscape where responsiveness is an emergent property of cell identity rather than a simple function of receptor abundance. Our results demonstrate that BDNF signaling is governed by a multi-layered gating system: (1) receptor levels initialize the potential for pathway induction, (2) a sustained reduction of surface TrkB serves as predictor of increased phosphorylation of downstream signaling effectors, and (3) cell-intrinsic environments act as the final arbiter of BDNF sensitivity. By moving beyond the ‘average’ signal of bulk biochemistry, we show that even identical receptor profiles yield distinct signaling profiles across cell types, reframing BDNF sensitivity as a form of prepared competence. This multi-layered regulation provides a framework for understanding how cellular context dictates the interpretation of the neurotrophic message.

## RESULTS

### Single-cell mapping resolves a continuous maturation gradient and lineage diversity

To determine how individual neural identities respond to a uniform BDNF stimulus, we utilized single-cell mass cytometry to simultaneously quantify 19 signaling effectors across primary spinal cord cultures (E14 rat) (Fig. 1A, table S1-2). Treatment with continued deprivation (ContDepr), reapplication of complete media (Rescue), BDNF, and co-treatment of BDNF with a pan-Trk inhibitor, K252A, enabled comparison of cell signaling responses across a time course (0, 5, 15min, 1, 4, 16hr). We selected E14 rat spinal cord as our model system because it provides a developmental window in which neural progenitors, neurons of varying maturity, and non-neuronal populations coexist within a single preparation. Thus, the heterogeneity of this culture system enabled our simultaneous assessment of BDNF signaling across multiple lineages and maturation states.

**Fig. 1.**
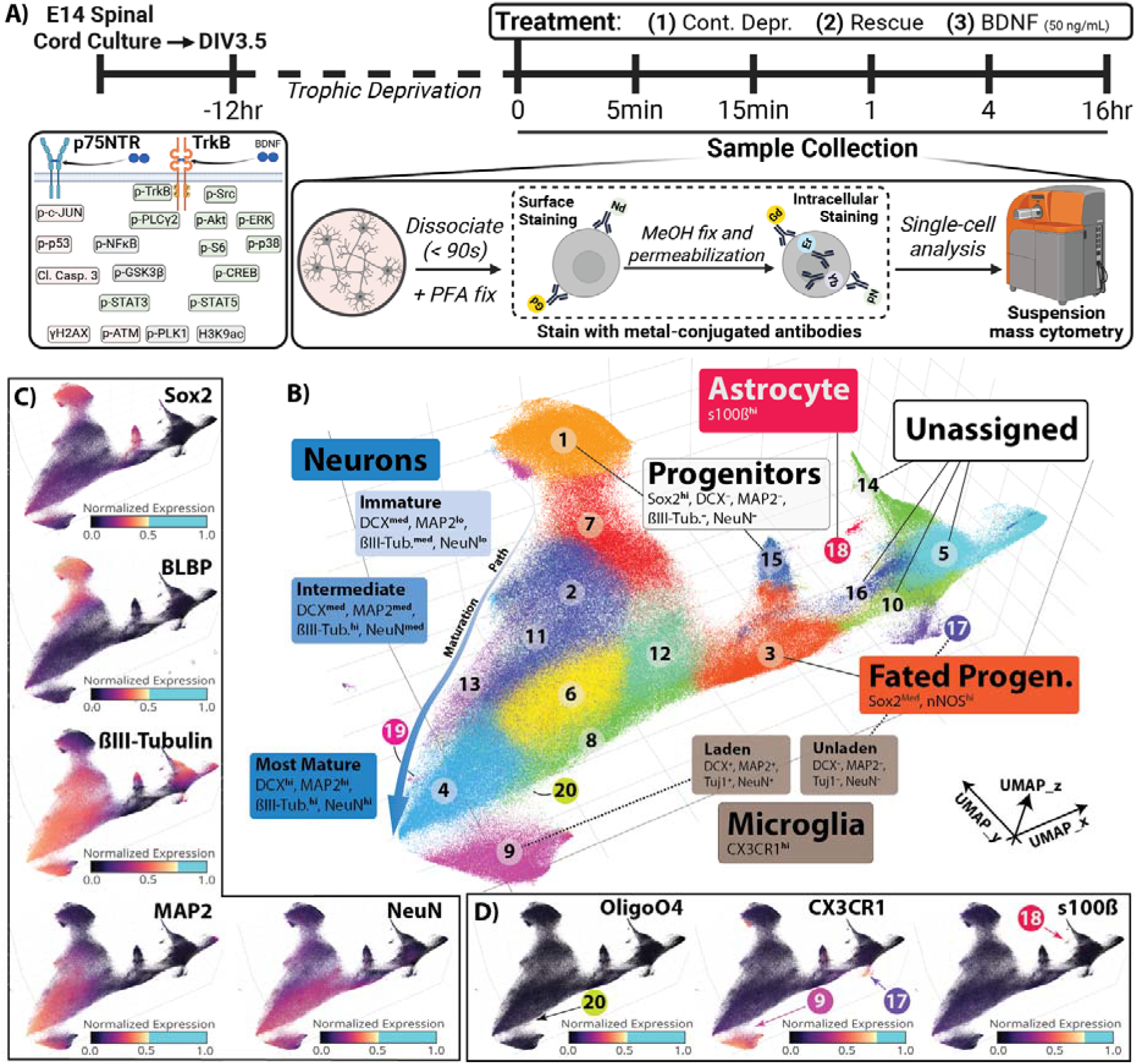
Single cell mass cytometry identifies distinct cell types in rat embryonic spinal cord cultures. (**A**) Embryonic day 14 (E14) rat spinal cord are cultured to day in vitro (DIV) 3.5 before a 12hr supplement starvation and subsequent treatment. Samples are collected at indicated time points by rapid dissociation (<90s) and fixation by 2% paraformaldehyde (PFA), then stored at -80°C. Samples are barcoded, pooled, and simultaneously stained with isotopically pure metal-conjugated antibodies before and after methanol (MeOH) permeabilization and fixation. Stained samples are then analyzed by single-cell (suspension) mass cytometry. Created using BioRender. (**B**) Leiden clustering by cell identity markers is visualized by dimensionality reduction on a 3-dimensional Uniform Manifold Approximation and Projection (UMAP) colored and labeled by identified cell types. (**C**) The resulting UMAP is colored by the level of individual cell identity markers to highlight the observed neuronal maturation gradient that shifts from Sox2+, BLBP+ progenitors to more mature neuronal identity with increasing levels of ßIII-Tubulin, MAP2, and NeuN. (**D**) Non-neuronal cell identity marker levels are plotted to define glial cell populations. Individual cell identity marker expression is normalized from 0 to 1, and gradient is shifted to emphasize locations of marker enrichment, with cells above the intensity threshold colored cyan.

Leiden clustering of identity marker expression distinguished 20 unique cell populations (Fig. 1B, fig. S1A), which were partitioned into four primary categories based on marker expression level (fig. S1A, B): Progenitor (Sox2+, 12.8%), Neuronal (ßIII-tubulin+, 52.5%), Non-Neuronal (17.0%), and Unassigned (17.6%). Unassigned cells represent those that could not be unambiguously defined by our panel of identity markers.

Notably, based on expression levels of neuronal identity markers, we were able to resolve a developmental trajectory originating from Sox2+ progenitors (Cluster 1), through intermediate immature neurons (Clusters 7, 2, 13), to the most mature NeuN-hi neurons (Cluster 4) (Fig. 1C, fig. S1D). We identified additional Non-Neuronal cell types, including nNOS+ fated progenitors, astrocytes (s100ß+), oligodendrocytes (OligoO4+) and microglia (CX3CR1+) based on their distinguishing identity marker (Fig. 1C, fig. S1A, B). Microglia can be further classified based on the presence (“laden”) or absence (“unladen”) of neuronal identity markers, which may represent the phagocytic state of these CX3CR1+ cells. By resolving this maturation gradient and the cell-type heterogeneity at single-cell resolution, we established a high-dimensional system to test to what extent BDNF signaling is a universal event or a state-dependent property of specific lineages.

### The global BDNF response is driven by a discrete subpopulations of cells

To determine if our platform recapitulated well-established canonical signaling profiles, we first aggregated our single-cell data into a “pseudobulk” format to assess the global response to BDNF (fig. S2-3). As expected, BDNF induced rapid activation across core Ras/MAPK and PI3K/Akt pathways, including pERK, pAkt, and pCREB, relative to controls where BDNF deprivation was continued (Fig. 2A, fig. S4A). These responses were largely abrogated by the Trk inhibitor K252a (Fig. 2B, fig. S4A). In addition, BDNF stimulation exhibited a unique phosphoproteomic profile which was distinct from the general trophic response elicited by refeeding with rich medium (“Rescue”) (fig. S5), confirming the specificity of the BDNF stimulus. However, single-cell analysis indicates that these aggregate trends obscure observable heterogeneous responses. At peak activation (15 min), not all cells respond uniformly, but instead discrete subpopulations contribute to the pseudobulk response, ranging from 47% to 75% of cells (Fig. 2C, fig. S4B, table S3). This observation provides evidence that BDNF responsiveness is not uniform but is restricted to specific cellular contexts (Fig. 2D).

**Fig. 2.**
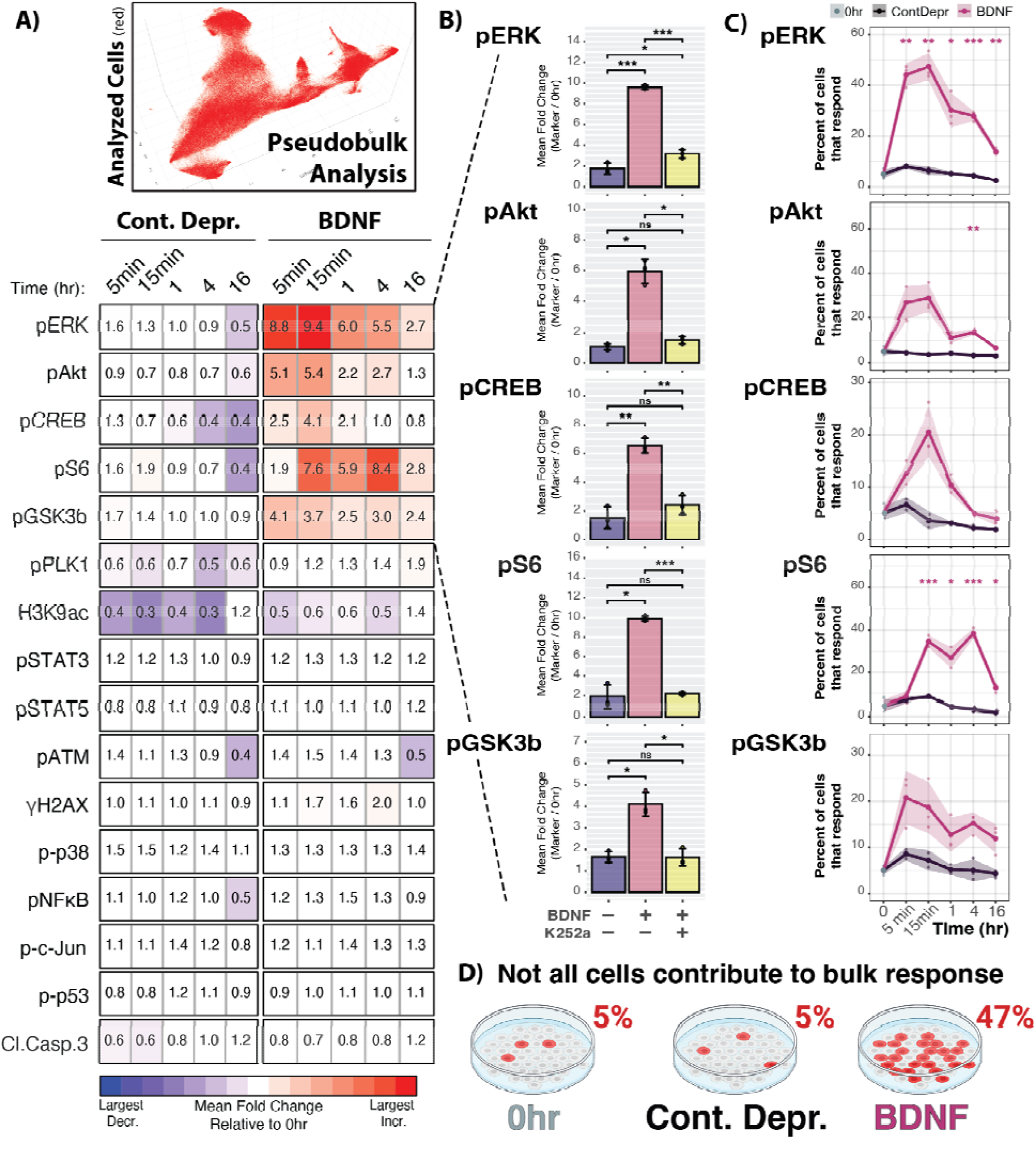
Pseudobulk analysis defines responsive subpopulation of cells responsible for global BDNF-induced signaling. (**A**) All cells were included in the pseudobulk analysis of the mean fold change (FC) of responsive cells relative to 0hr, as depicted by heatmap for select markers across time after either continued deprivation (Cont. Depr.) or BDNF treatment. No change is indicated by white (0.67 < FC < 1.5), the level of decrease is visualized by a blue scale (FC ≤ 0.67), and the level of increase is visualized by a red scale (FC ≥ 1.5). Values represent the mean FC (0hr, n = 4; Cont. Depr., n = 2-3; BDNF, n = 3). (**B**) Co-treatment of BDNF with a Trk inhibitor (K252A) largely abrogated BDNF-induced responses at 1hr of treatment. Points represent mean±SD of 3 culture wells (n = 3) from 1 litter. (**C**) The percent of cells that are defined as responsive plotted for individual signaling markers across time. Points represent mean±SD of 2-4 culture wells (0hr, n = 4; Cont. Depr., n = 2-3; BDNF, n = 3) from 2-3 litters. (**D**) Schematic visualization depicting the cells that contribute to a bulk readout, where not all treated cells respond to the stimulus. Created in BioRender. (B,C) p-values are adjusted for multiple comparisons (Welch’s unpaired two-sided t-tests with Holm adjustment): * p < 0.05, ** < 0.01, *** < 0.001; ns = not significant. (C) Welch’s ANOVA across time points within each treatment give p ≤ 0.006 for all five markers after BDNF (pERK and pS6 < 0.001) and p ≤ 0.011 for pERK, pS6, and pCREB in ContDepr (p > 0.25 for pAKt and pGSK3b). See Supplemental Table 3 for values.

### Maturation state and cell identity regulate BDNF signaling competency

We next determined if the heterogeneity of BDNF responsiveness was structured by cell identity, for instance if neuronal cells responded and non-neuronal cells did not. While both neuronal and non-neuronal cells show increased activation of established pathways (fig. S6B), neuronal cells exhibit a greater magnitude of change (fig. S6B). Inhibition of Trk activity with K252a significantly reduces BDNF-induced activation to control or near-control levels in both cell type categories (fig. S6C).

A more granular assessment of cell-type-specific pathway activation revealed a diversity of responsiveness (Fig. 3A-B, fig. S7A). Intriguingly, we observed a maturation-dependent acquisition of signaling competence along the neuronal lineage. Sox2+ progenitors remained relatively signaling-silent (1.7-fold pERK) compared to intermediate and mature neurons, which exhibited a 16.2- to 17.2-fold induction at the same time point (Fig. 3C, table S5).

**Fig. 3.**
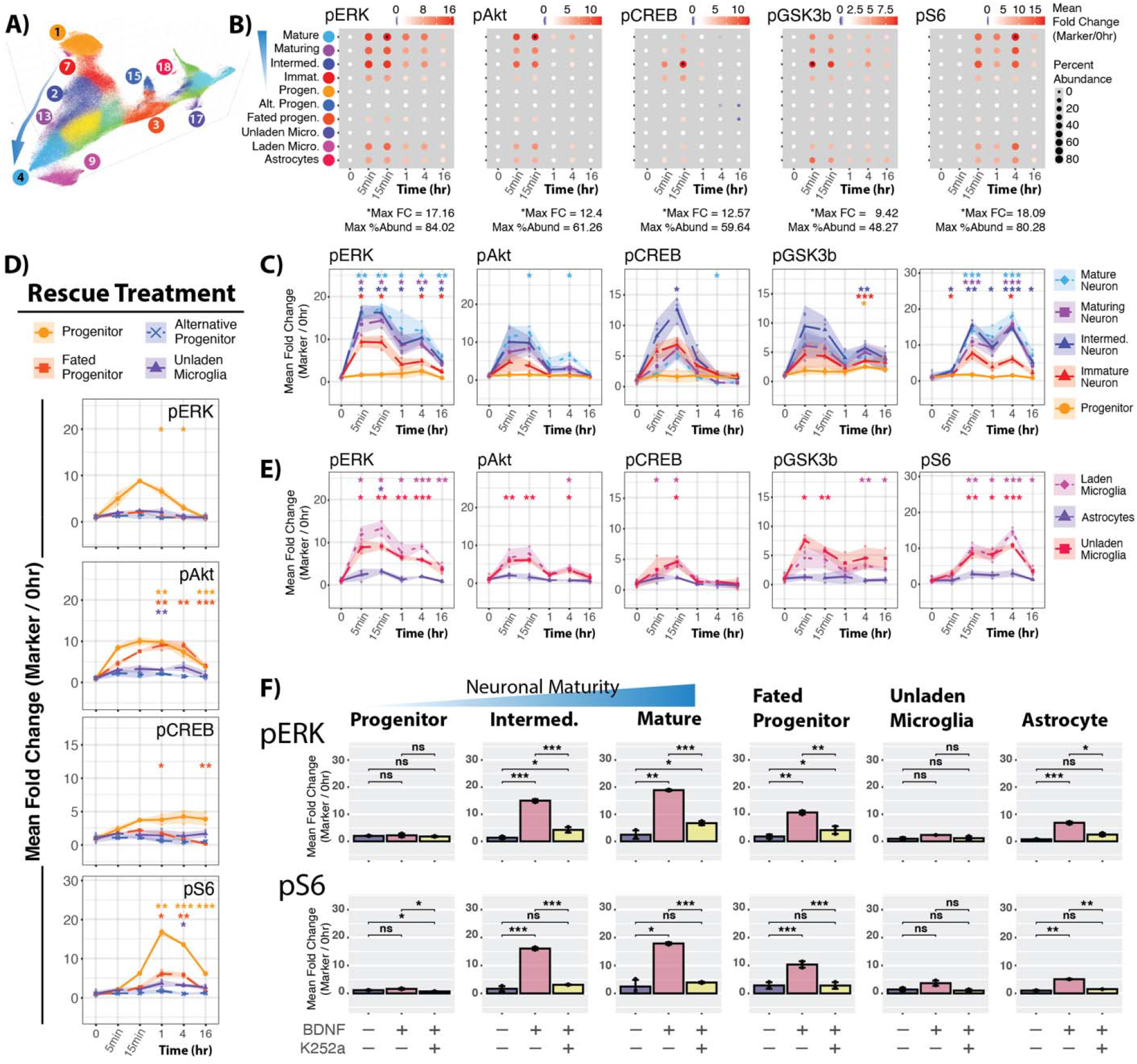
BDNF-induced signaling dynamics differ between identified cell types. (**A**) UMAP highlighting cell identity clusters that are subsequently analyzed. (**B**) Dot plot depicting the percent of cells that increase activation of the selected signaling molecule within each identified cell type. Size indicates the percent of the given cell type that contribute to the response, while color indicates the mean fold change relative to 0hr. (**C**) Line graph representation of select data from (B) emphasizes the comparison of responsive cells in the neuronal maturation path. (**D**) Signaling responses to complete-medium rescue treatment in cell types that do not exhibit a robust response to BDNF. (**E**) Line graph representation of select data from (B) to compare responses in non-neuronal cell types. (C-E) Points represent mean±SD of 2-4 culture wells (0hr, n = 4; Cont. Depr., n = 2-3; BDNF, n = 3) from 2-3 litters. (**F**) Co-treatment of BDNF with a Trk inhibitor (K252A) largely abrogated cell-type-specific BDNF-induced responses at 1hr of treatment Points represent mean±SD of 3 culture wells (n = 3) from 1 litter. (C-F) p-values are adjusted for multiple comparisons (Welch’s unpaired two-sided t-tests with Holm adjustment): * p < 0.05, ** < 0.01, *** < 0.001; ns = not significant - see Supplemental Table 5 for values.

Crucially, progenitors showed robust activation when complete-medium was reapplied (Fig. 3D, table S5), confirming that they are intrinsically signaling-competent but specifically insensitive to BDNF. Further comparison of treatment-dependent responses shows a relatively consistent response to complete-medium rescue along the neuronal lineage, suggesting that maturation-dependent responsiveness is not generalizable to any stimulation (fig. S8).

Non-neuronal lineages similarly showed divergent sensitivity: astrocytes and “laden” microglia were highly responsive, whereas “unladen” microglia remained signaling-silent (Fig. 3E). Furthermore, the Trk inhibitor K252a reduced most BDNF-induced responses to control or near-control levels (Fig. 3F, fig. S7B, table S5). pERK was not fully reduced to control levels in intermediate and mature neurons and fated progenitors (Fig. 3F), nor was pAkt in fated progenitors and intermediate neurons (fig. S7B). Together, these data suggest that BDNF induces cell type-specific responses that are largely sensitive to Trk inhibition.

### Unsupervised clustering reveals cell identity-dependent BDNF signaling signatures

By measuring 19 signaling markers at once, we were able to define signaling patterns in identified cell types based on the temporal changes of individual marker abundance. However, assessing each marker separately limits our analysis and does not take advantage of our multi-dimensional dataset. Thus, before exploring the observation of Trk-activity dependence, we expanded on our characterization of responses to BDNF by defining “signaling signatures” based on input from all measured signaling markers without regard to cell type.

To move beyond separate analysis of individual markers, we performed unsupervised Leiden clustering based exclusively on signaling marker abundance, agnostic to cell identity (Fig. 4A). This re-clustering analysis resolved 21 signaling clusters (fig. S9A) distinguished by their marker expression profiles (fig. S9B). By plotting cells back onto the generated signaling UMAP by time and treatment, we were able to observe shifts in the distribution of cells to define signaling signatures. Based on the shift in distribution upon BDNF stimulation (Fig. 4B), we identified three temporal signaling signatures – Signature A (rapid pAkt/pERK), Signature B (sustained pS6), and Signature C (complex multi-marker activation) – as well as Basal and Non-Signaling Signatures (Fig. 4C). Of the 21 signaling clusters identified by the clustering analysis, 13 were represented in these defined signaling signatures, each exhibiting a unique signaling marker expression profile (Fig. 4D).

**Fig. 4.**
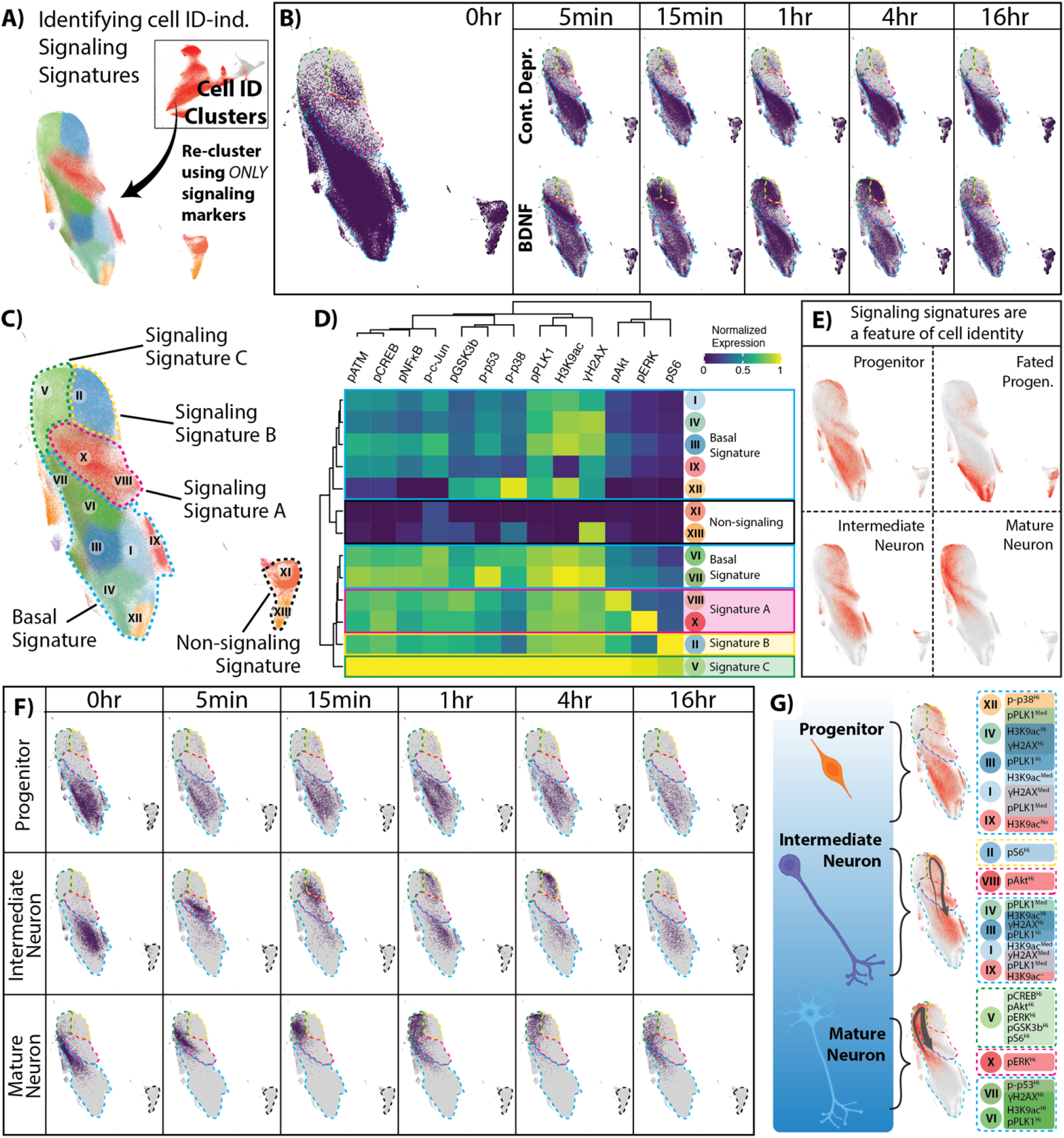
BDNF induces signaling signature defined by time and by cell type. (**A**) Depiction of re-clustering of cell ID clusters (excluding Unassigned) using signaling markers to identify signaling signatures that are agnostic to the initial cell identity. (**B**) UMAP colored by time point within continued deprivation or BDNF treatment to show the temporal shift in cell distribution across the signaling signature landscape. (**C**) UMAP of signaling clusters that dynamic shift in cell distribution upon BDNF induction. Signaling clusters are categorized into Non-signaling Signature, Basal Signature, Signaling Signature A, Signaling Signature B, and Signaling Signature C based on the BDNF-induced temporal shifts in distribution. (**D**) Expression heatmap for indicated signaling clusters to molecularly define the identified signaling signatures. (**E**) Select cell ID clusters are overlayed on the signaling signature UMAP to identify signaling-driven grouping of cells. (**F**) UMAP colored by time point across cell types treated with BDNF to depict temporal shift in cell distribution. (**G**) Signaling dynamics are summarized for selected cell types. Created in BioRender.

Surprisingly, when we mapped cell types back onto the signaling UMAP, they grouped together, even without identity markers in the input data (Fig. 4E). When we plotted the distribution of individual cell types over time, each followed a unique temporal profile. Intermediate neurons progressed primarily through Signature B, whereas mature neurons adopted the more complex Signature C (Fig. 4F). Although these temporal snapshots represent population shifts rather than individual cell trajectories, the divergence between mature and intermediate neurons shows that maturation state defines not just the magnitude, but the “signature” of BDNF signal induction. These data suggest that BDNF responsiveness is a programmed feature of cell identity, where different lineages decode the same ligand into unique, temporally regulated signaling signatures (Fig. 4G).

### The sustained reduction of surface TrkB correlates with BDNF sensitivity

We next sought to identify the molecular mechanisms governing these distinct signaling signatures. A primary candidate for this variability is the relative abundance of the two BDNF receptors themselves; however, it remains unknown to what degree TrkB and p75NTR density dictate signaling magnitude and pathway specificity across cell populations.

To resolve this, we first quantified the contribution of total TrkB receptor expression using a tiered pseudobulk analysis, where each cell was categorized based on its total level of TrkB (Fig. 5A; “None”, “Lo”, “Med”, “Hi”). While 91.23% of cells express some level of TrkB, receptor density was not a consistent predictor of signaling output (Fig. 5B, fig. S10A). Increased activation of effectors downstream of BDNF-TrkB, such as pERK and pS6, were primarily associated with TrkB presence; however, pCREB was not directly correlated with increased TrkB levels, suggesting TrkB abundance is not the only factor driving its activation (Fig. 5B). Cells not expressing TrkB (“None”) showed no activation of certain signaling markers, such as pNF-kappa-B and p-c-Jun, though this population of cells did exhibit a modest increase in pERK, pS6 and pCREB (Fig. 5B). The increase in pERK in the “None” cell population aligns with our observation that Trk inhibition only partially suppresses ERK activation (Fig. 2C), which may suggest Trk-independent activity.

**Fig. 5.**
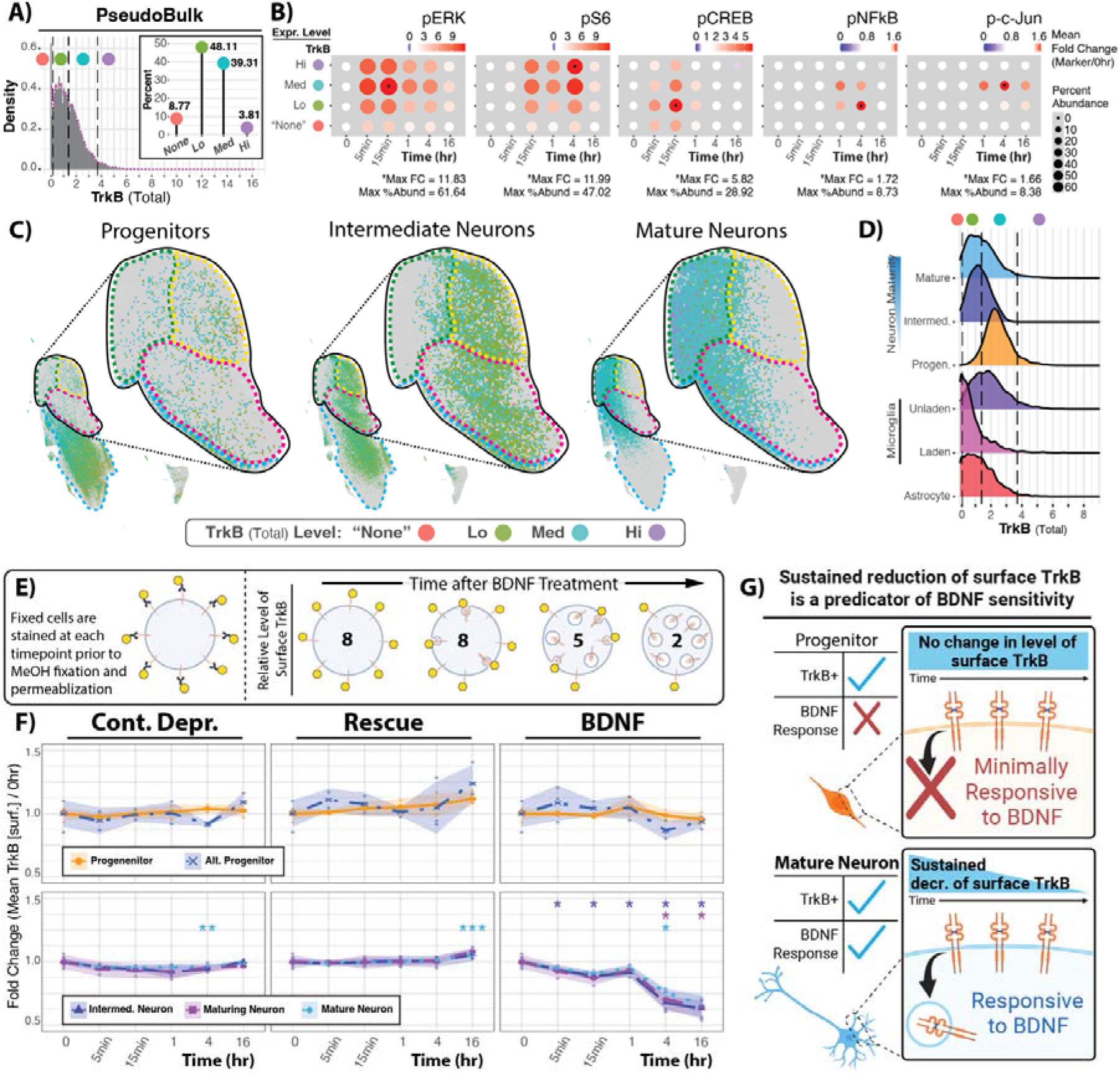
BDNF-responsive cell types exhibit sustained reduction of surface TrkB. (**A**) Distribution and categorization of TrkB (total) expression level in Pseudobulk data. Inset depicts percent of all cells within each expression category. (**B**) Pseudobulk signaling response within each expression category to define general trends correlating to TrkB expression. (**C**) Distribution of signaling signatures by cell ID and colored by TrkB (total) expression levels. Dotted lines indicate Signaling Signature A (rapid pAkt/pERK; magenta), B (sustained pS6; yellow), and C (complex multi-marker activation; green). Insets highlight the BDNF-induced signatures. (**D**) TrkB (total) expression distribution across selected cell types, indicating each cell type expresses some degree of TrkB. Dashed lines represent the same values set on pseudobulk data in (A). (**E**) Diagram of TrkB surface staining, used to measure the reduction of surface TrkB as a correlate of internalization. (**F**) Fold change of mean TrkB (surface) relative to 0hr in response to continued deprivation (Cont. Depr.), Rescue, or BDNF within select cell types. Points represent mean±SD of 2-4 culture wells (n) from 2-3 litters. p-values are adjusted for multiple comparisons (Welch’s unpaired two-sided t-tests with Holm adjustment): * p < 0.05, ** < 0.01, *** < 0.001; ns = not significant. See Supplemental Table 7 for values. (**G**) Results suggest that the reduction of surface TrkB predicts BDNF responsiveness better than total TrkB abundance. Created in BioRender.

To assess whether different levels of TrkB influenced signaling profiles in different cell types, we incorporated TrkB expression level categories into our signaling signature analysis (Fig. 5C). In cell types with robust responses, such as intermediate and mature neurons, all levels of TrkB in the given cell type were present in BDNF-responsive (insets; Signatures A-C) and Basal Signatures (blue dashed outline). Within signaling signatures, there is some grouping based on TrkB level – most notably, TrkB-Hi in mature neurons in Signature C (green dashed outline) – suggesting some contribution of TrkB level to the temporal signaling profile. Although fewer in number, some progenitor cells were present in BDNF-responsive signatures (inset), which had varying levels of TrkB. However, the majority of progenitors exhibit a Basal Signature (blue dashed outline in Fig. 5C), despite harboring Med-to-Hi levels of TrkB (Fig. 5D). This presents an apparent paradox whereby TrkB-expressing cells (i.e. progenitors) can be insensitive to BDNF.

One possible mechanism for the TrkB-high, signaling-silent progenitors is that TrkB in progenitors might not undergo endocytosis, a process thought to be required for BDNF signaling. Using a surface-specific staining protocol, we quantified ligand-induced TrkB depletion from the plasma membrane (Fig. 5E). Our results show a clear correlation between signaling competence and a reduction of surface receptor abundance (Fig. 5F). While intermediate and mature neurons showed significant surface TrkB depletion at later time points, unresponsive progenitors exhibited stable surface profiles. This lack of surface TrkB reduction provides a correlative explanation for why receptor-positive cells remain functionally signaling-silent. These data indicate that the reduction of surface receptors over time is an important part of BDNF regulation (Fig. 5G), suggesting that surface receptor dynamics may serve as a predictor of responsiveness beyond static protein abundance alone.

To assess the contribution of p75NTR expression on BDNF-induced signaling, we performed a tiered pseudobulk analysis based on the level of p75NTR in each cell. Comparison of expression level categories shows an apparent shift in signaling patterns: high p75NTR density was associated with increased stress-responsive p-c-Jun and p38 activation, but also unexpectedly amplified trophic markers like pCREB and pERK (fig. S10B). Taken together, these analyses suggest that receptor presence alone, regardless of density or receptor profile, is insufficient to confer signaling competence, which is predicted by the sustained time-dependent reduction of surface TrkB.

### Cell identity modulates signaling response set by receptor stoichiometry

Our results show that the abundance of each BDNF receptor individually contributes to the BDNF response but does not account for all observed signaling changes. This is expected, as others have established that interplay between TrkB and p75NTR can tune signaling responses (*7–11*). We therefore wanted to understand how receptor stoichiometry influences the profile of BDNF-induced signaling.

To address this question, we first compared signaling responses in cells with different combinations of TrkB and p75NTR, regardless of cell identity. By plotting the level of p75NTR vs total TrkB expression, we segmented four categories of cells based on our previously-applied tiered pseudobulk analysis: p75NTR–/TrkB–, p75NTR+/TrkB–, p75NTR–/TrkB+ and p75NTR+/TrkB+ (Fig. 6A). Consistent with synergistic models, we observed that TrkB and p75NTR act in concert to amplify markers like pERK, pS6, pCREB, and pNF-kappa-B, while p75NTR expression alone – in the absence of TrkB – was sufficient to drive p-c-Jun activity (Fig. 6B, fig. S11).

**Fig. 6.**
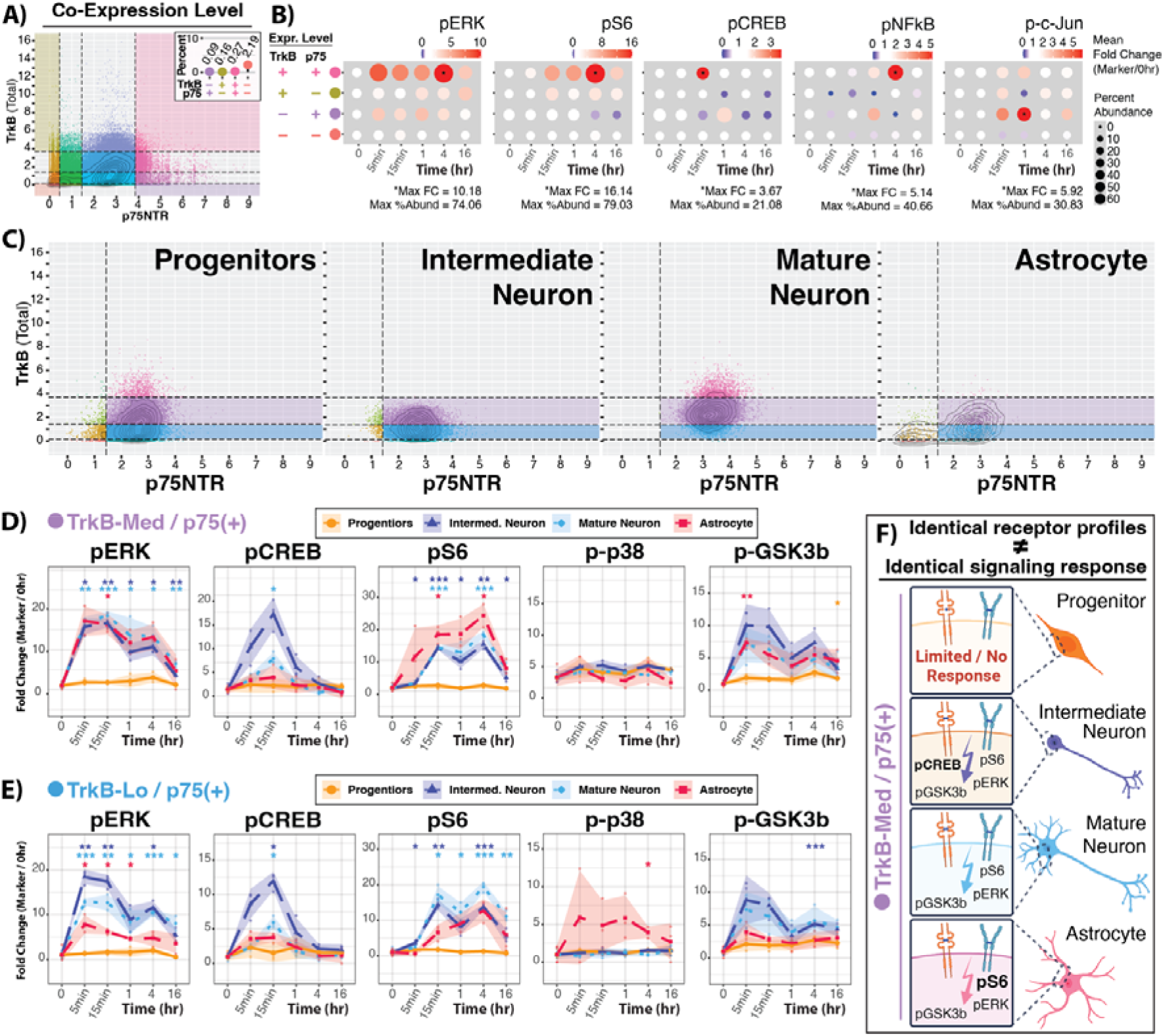
Identical BDNF receptor profiles do not confer identical responses across cell types. (**A**) Distribution and categorization of TrkB and p75NTR expression level in Pseudobulk data. Inset depicts percent of all cells within select expression categories defining TrkB–/p75–, TrkB–/p75+. TrkB+/p75–, TrkB+/p75+ subpopulations. (**B**) Pseudobulk signaling response within each expression category highlighted in (A) to define general trends correlating to TrkB, p75NTR co-/expression. (**C**) Distribution and categorization previously characterized cell types based on TrkB, p75NTR co-/expression. (**D, E**) Comparison of signaling dynamics across cell types with a (D) TrkB-Med/p75+ or (E) TrkB-Lo/p75+ expression pattern. (**F**) Schematic depicting cells with the same BDNF receptor profile exhibiting different signaling responses. Created in BioRender. (D-E) Points represent mean±SD of 2-4 culture wells (0hr, n = 4; Cont. Depr., n = 2-3; BDNF, n = 3) from 2-3 litters. p-values are adjusted for multiple comparisons (Welch’s unpaired two-sided t-tests with Holm adjustment): * p < 0.05, ** < 0.01, *** < 0.001; ns = not significant. See Supplemental Table 8 for values.

We next asked how receptor expression levels influence BDNF-induced signaling responses within a given cell type. Using astrocytes as an example, we observe synergistic amplification of certain signaling markers, where p75NTR+/TrkB+ astrocytes exhibited high activation of pERK and pS6 with a milder increase in pAkt (fig. S12). Other signaling marker activity did not show synergy: pGSK3b exhibited similar responses in TrkB-med cells regardless of p75NTR presence, and p-p53 was only increased in p75NTR–/ TrkB-med, suggesting a TrkB-dependent/p75NTR-independent response. Together, this analysis shows that receptor stoichiometry can influence BDNF-induced signaling in the same cell type.

To determine if receptor expression level is the ultimate determinant of signaling output, we then analyzed signaling responses across different cell types, holding the receptor expression profile constant. We isolated subpopulations of astrocytes, progenitors, intermediate neurons, and mature neurons that were matched for identical surface levels of both TrkB and p75NTR (Fig. 6C). Surprisingly, an identical receptor profile (for instance TrkB-med/p75(+)) did not result in an identical response. Instead, each cell type produced distinct signaling patterns (Fig. 6D). This observation held when comparing cell-type specific responses in a different receptor profile (TrkB-lo/p75(+); Fig. 6E, fig. S13). Additional comparison of the same cell type across these two receptor profiles highlights our observation that receptor stoichiometry can modulate signaling responses.

These results ultimately demonstrate that while receptors are necessary for BDNF responsiveness, they are not sufficient to dictate the signaling pattern (Fig. 6F). Instead, the receptor stoichiometry sets the potential of BDNF sensitivity, which is further tuned by lineage-dependent regulators. These observations suggest that BDNF signaling is a hierarchical process whereby the cell’s internal state determines the signaling outcome of this single extracellular cue.

## DISCUSSION

The signaling response to BDNF is traditionally understood through receptor biology. Our results support a more holistic model in which receptor stoichiometry sets the potential for BDNF sensitivity, and the intrinsic environment allows for a cell to engage that potential. By moving beyond the “average” signals of bulk biochemistry, our high-dimensional analysis reveals a BDNF-induced signaling landscape where identical receptor profiles produce differing signaling patterns across various cell types. By reframing BDNF sensitivity as a form of prepared competence, we can view BDNF-induced signaling as interconnected points of regulation (Fig. 7). While receptor stoichiometry primes a cell to respond to BDNF, additional factors that are intrinsic to individual cell types – such as surface receptor dynamics – determine exactly when and how the BDNF message is translated into a functional outcome.

**Fig. 7.**
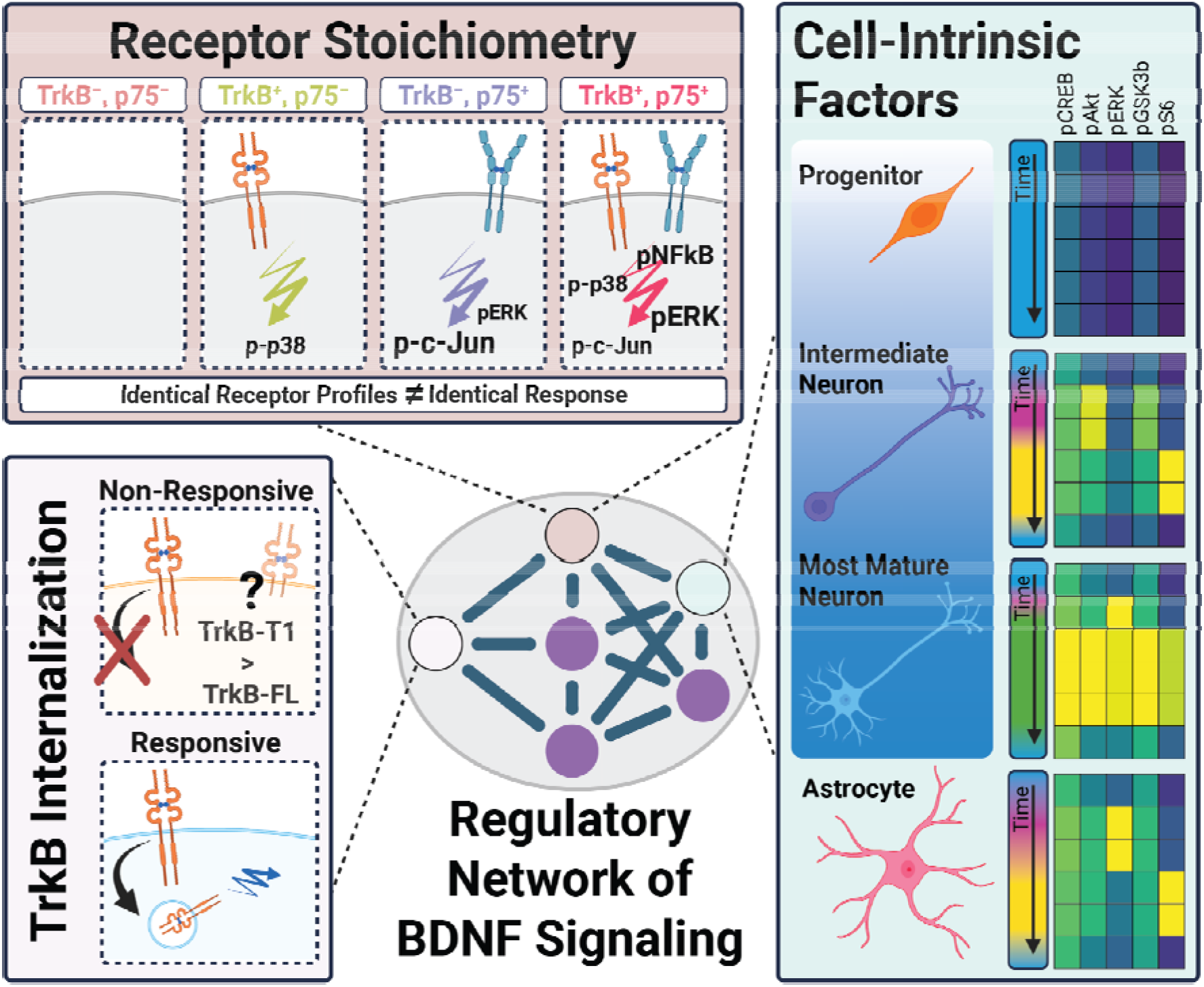
Factors contributing to the heterogeneity of BDNF signaling responses. Stimulation with BDNF induces different cell signaling responses across different cell types. The pattern of BDNF receptor abundance is necessary to regulate how cells respond; however, stoichiometry alone is not sufficient to explain how some cell types are non-responsive, despite high levels of TrkB. A better predictor of BDNF sensitivity is the prolonged reduction of surface TrkB, a correlate of internalization, where the ratio of TrkB isoforms may add a level of developmental competency to the overall regulation network. Ultimately, the cell-intrinsic environment acts as the final filter through which a cell interprets the BDNF message. Created in BioRender.

### Cell type-specific context defines BDNF response set by receptor stoichiometry

A prevailing assumption of receptor-ligand signaling posits that receptor ratio and density completely predict signaling outcomes. However, by isolating subpopulations with matched receptor expression patterns across unique lineages, we demonstrate that this model is incomplete. To our surprise, identical receptor profiles yielded largely different signaling programs across cell types: a mature neuron’s response is fundamentally different from that of an astrocyte or progenitor even at the same TrkB/p75 stoichiometries. This indicates that the interpretation of a BDNF signal is not invariantly set by the receptor stoichiometry itself but is modulated by the intracellular context. Our results expand upon computational models of NGF/TrkA signaling which suggest that protein expression noise – specifically intrinsic noise (*15, 16*) – drives single-cell ERK variability (*12, 17*). Furthermore, our multiplexed measurements provide a concrete example of a biological context that sets the ‘cell state’ described in these models: the extent of neuronal differentiation sets a maturation gradient with distinct BDNF responsiveness. This suggests that previously observed stochastic variance may actually reflect programmed, deterministic differences across neuronal maturity, thus reframing ‘noise’ as a developmentally regulated feature.

### Sustained reduction of surface TrkB is a functional correlate of responsiveness

If receptor density cannot predict signaling output, does receptor trafficking serve as the primary indicator of responsiveness (*18*)? Following ligand binding, TrkB is internalized into signaling endosomes to trigger canonical trophic cascades (*6, 19*). Our data indicate a correlation between signaling competence and receptor trafficking: while neurons and astrocytes show significant ligand-induced surface TrkB depletion, non-responsive populations – most notably progenitors – exhibit stable surface receptor levels. Similar to other signaling systems modulated by receptor internalization (*20*), our observations suggest a trafficking-based explanation for why receptor-positive cells remain signaling-silent. While these findings are correlative and do not yet prove a causal requirement for internalization, the consistency of this pattern across neural lineages suggests that trafficking kinetics are a more reliable predictor of BDNF sensitivity than static protein abundance. Currently, the mass cytometry platform cannot measure internalization directly, and different tools are needed to test these ideas decisively. Defining the precise molecular regulators of this internalization at single-cell resolution remains a key open question to be addressed by future endocytic perturbation studies.

Furthermore, the “progenitor paradox” (i.e. high TrkB expression paired with a limited reduction in surface TrkB) may represent a point of signaling regulation rather than an inability to respond. In this context, progenitor non-responsiveness could be a mechanism of “prepared competence,” where cells maintain a high TrkB density as they await a developmental window that permits differentiation. A compelling candidate for this gate is a developmental switch from truncated (TrkB-T1) to full-length (TrkB-FL) isoform expression. Under this hypothesis, TrkB-T1 would act as a signaling sink or dominant-negative regulator (*21, 22*) until TrkB-FL expression becomes dominant, which would have profound consequences for progenitor cell-fate decisions (*23, 24*). Although our antibody targets the shared extracellular domain and cannot discern these isoforms, our data are consistent with this hypothesized cell-intrinsic factor: the presence of the receptor but a lack of sustained surface reduction and signaling response. Alternatively, the time points for this study may not capture rapid (< 5 min), transient TrkB-dependent activity at the cell surface (*25*). Thus, further investigation is warranted into this potential mechanism of prepared competence, and confirmation of this “active gate” will require genetic perturbation, more refined time courses, and isoform-specific proteomic probes.

### Receptors are necessary, but not sufficient in BDNF signaling regulation

While receptor trafficking provides a rationalization of BDNF responsiveness, how does the interplay between TrkB and p75NTR shape BDNF signaling? Our results suggest that BDNF pathways utilize a combinatorial logic where the level of receptor availability tunes specific cascades. Consistent with established synergistic models (*7, 9, 22, 23*), we observe that TrkB and p75NTR act synergistically to amplify trophic pathways like pERK and pNF-kappa-B, while p75NTR alone is sufficient to drive stress-responsive p-c-Jun activity. We posit that these patterns may be further regulated by proteolytic cleavage of p75NTR, which releases its intracellular domain (24), adding another layer of receptor-mediated regulation. Our high-resolution mapping highlights that p75NTR is not merely a “death receptor” but a critical synergistic partner that heightens cellular sensitivity to the BDNF message in the context of our experimental system.

It is unsurprising that receptors are necessary to initiate BDNF-induced signaling; however, our results indicate that receptor presence alone is not sufficient to dictate the final output. We demonstrate that this receptor interplay is modulated by cellular context unique to each lineage. This is evidenced by our finding that equivalent receptor expression patterns yield varying signaling programs (Fig. 6), that only a discrete subpopulation (∼47–75%) of total cells respond to BDNF stimulation (Fig. 2), and that unsupervised clustering based solely on signaling markers independently groups cells by their biological identity (Fig. 2).

### Intrinsic factors ultimately gate BDNF interpretation across distinct cell types

While we cannot determine from fixed data whether individual cells traverse signaling signatures sequentially or preferentially occupy different signatures, insights from these data – made possible through high-dimensional single-cell proteomics – suggest a hierarchy of filters. These intrinsic filters may include receptor isoform ratio (*21, 22*), cleavage and internalization dynamics (*24*), cell cycle phase (*26*), or the epigenetic landscape (*27–29*), which likely determines the accessibility of target genes for BDNF-induced expression. Such multifactorial logic – where the signaling pattern rather than amplitude encodes biological meaning – mirrors findings in other systems, including calcium oscillations (*30–33*), TNF-NF-kappa-B dynamics (*34–36*), Notch signaling (*37*), growth factor-induced PI3K activity (*38*), and Wnt signaling (*39, 40*). By demonstrating that lineage modulates stoichiometry, we reframe BDNF signaling as a tiered process where maturation state defines the functional meaning of the extracellular cue.

### Current limitations and future study

While our high-dimensional approach provides additional insight into the regulation of BDNF signaling, several limitations of the current study warrant consideration. (**1**) **Limited antibody panel.** Although suspension mass cytometry enables simultaneous readout of multiple parameters at single cell resolution, the antibody panel limits which parameters are measured. Selected cell identity antibodies enabled distinction of unique cell types and maturation stages; however, cell identity definitions are descriptive and may be reductive. For example, astrocytes are defined by high s100ß abundance relative to other identity clusters, but maturity of these cells cannot be assessed. Astrocyte maturation could contribute to BDNF-induced signaling responses, since expression of TrkB isoforms is dependent on maturity (TrkB-FL to TrkB-T1) (*41*). The selected antibody panel also did not include other cell surface receptors, such as other Trk receptors or EGFR (*42*), which could lead to transactivation of observed Trk-independent signaling responses (Fig. 2B, 5B). Future studies can ask whether there is interplay between TrkB and other receptors, beyond p75NTR. In our hands, the phospho-TrkB (Y816) antibody did not reliably report BDNF stimulation in this assay, so we draw no conclusions from that channel; antibodies against the activation-loop tyrosines were not available validated for mass cytometry, but would give a more direct measure of TrkB activity. (**2**) **K252A is a broad kinase inhibitor.** Although commonly used as a pan-Trk inhibitor, K252A also inhibits other kinases (*43*), including PKA, PKC and PKG at nanomolar concentrations (*44*), and it cannot distinguish TrkB-from TrkA- or TrkC-dependent responses. Our study uses this compound to suggest TrkB-dependency of BDNF-induced responses, but did not include a K252A-alone control to prioritize time course samples, and the 1hr K252A experiment used cultures from a single litter (table S2). Because K252A inhibits not just Trk receptors, the responses it reduces may not all be Trk-dependent. K252A co-treatment reduced most BDNF-induced responses to near-control levels, except for pERK in pseudobulk (Fig. 2B) and in select cell types (Fig. 3F, fig. S7B). Where pERK persisted under K252A, it remained 2.5- to 6.7-fold above 0hr, compared with 0.7- to 2.5-fold under continued deprivation (tables S3 and S5). Off-target inhibition would be expected to lower, not preserve, signaling, so this residual activity more likely reflects incomplete Trk inhibition at 1hr or Trk-independent activation of ERK. Consistent with the latter, we observe pERK activity in cells with TrkB below our expression cutoff (TrkB “None”; Fig. 5B), although these cells may retain low TrkB levels, and in TrkB–/p75+ cells (Fig. 6B), suggesting possible routes through p75NTR or receptor transactivation. Orthogonal approaches, such as genetic loss of TrkB or a TrkB-selective antagonist like ANA-12 (*45*), should be considered in future experimental designs. (**3**) **Correlation of reduced surface TrkB with signaling.** Our findings regarding the reduction of surface TrkB and downstream signaling are correlative. Although the lack of receptor trafficking in progenitors serves as a consistent predictor of signaling silence, future investigation is required to establish a direct causal requirement for internalization in this specific context. (**4**) **Experimental paradigm.** Our analysis relies on discrete temporal snapshots. While these timepoints capture the evolution of signaling signatures across the population, they cannot directly track the dynamic trajectories of individual cells as they transition between signaling states. Additionally, the 12hr deprivation prior to treatment potentially limits the generalizability of our results. Our data show that trophic deprivation per se influences cell signaling: the continued deprivation time course alters pERK, pS6, and pCREB (Fig2C). Thus, conclusions regarding signaling competency are limited to this experimental set up until they are orthogonally supported. Alternatives or additions to the deprivation approach should be considered in extensions of this work. Furthermore, our findings are derived from an in vitro primary culture system; while this allowed for the high-resolution mapping of endogenous spinal cord lineages, future studies are required to determine how these nodes of regulation are modulated within the complex three-dimensional environment of the in vivo niche. Our high-dimensional dataset enables hypothesis generation; however, to establish functional significance of the observed signaling profiles, our generated hypotheses will require readouts of downstream consequences, such as cell proliferation by BrdU imaging or transcriptional changes by cell sorting and RT-qPCR. Finally, our study exclusively examines the response to mature BDNF. Because proBDNF preferentially engages p75NTR rather than TrkB (*46–48*), comparing the signaling landscapes elicited by each ligand form would directly test whether the regulatory nodes we describe are reconfigured when the receptor entry point shifts from TrkB to p75NTR. This could potentially identify an additional layer of ligand-form-dependent route diversity.

Beyond fundamental signaling, these findings suggest a new logic for neurotrophic drug development for neurological diseases and disorders. Our model predicts that global TrkB agonists will exhibit cell-type-specific efficacy in vivo, potentially explaining the inconsistent results and partial success of BDNF-based therapies in clinical trials (*49, 50*). Effective strategies must likely move beyond simple receptor agonism toward combination approaches that shift the intracellular context – transitioning a cell from a signaling-silent state to a BDNF-sensitive one. For example, the use of maturation-stage-specific HDAC inhibitors could remodel the epigenetic landscape of stagnant populations (*27, 51*), lowering the “intrinsic filter” to permit TrkB-mediated differentiation. However, the transition in vivo faces the formidable challenge of cell-type-specific delivery (*50*), as ensuring context-shifting agents reach the intended subpopulation without inducing off-target effects in already-responsive neurons remains a hurdle. Ultimately, this study suggests that neurotrophin sensitivity is a form of prepared competence that lies not just in the ligand and the receptors, but in the internal machinery of the cell itself.

## MATERIALS AND METHODS

### 1. Animals

All experiments were conducted in accordance with AAALAC guidelines on a protocol approved by the University of Virginia ACUC. Timed pregnant Sprague Dawley rats were purchased from Charles River (Strain Code: 001) at embryonic day 12 (E12) and housed on a 12hr light/dark cycle prior to embryonic tissue dissection.

### 2. Cell culture

Primary dissociated neural cultures were prepared from E14 Sprague Dawley rat spinal cords. Briefly, spinal cords (sacral level excluded) were dissected in chilled HBSS, minced, and enzymatically digested for 15 min at 37°C in a filtered enzyme solution containing 0.25% (w/v) trypsin, 0.04% (w/v) hyaluronidase, and 0.04% (w/v) DNase. Tissue was mechanically dissociated by trituration in plating media (Basal Medium Eagle supplemented with 5% FBS, 1X N2 Supplement, 1X GlutaMAX, and 1% Penicillin-Streptomycin) and passed through a 70 µm cell strainer. Cells were plated at a density of 5x10^5^ cells/well in 12-well plates pre-coated with 0.1 mg/mL poly-D-lysine (overnight) and 15 µg/mL laminin (>2hr) in PBS. Initial cell viability was >90% as determined by Trypan blue exclusion prior to plating. Within 16hrs, plating media was replaced with Maintenance Media (Neurobasal media supplemented with 1X B27 and 1X GlutaMAX). Cultures were maintained at 37°C in 5% CO2, with half-volume media exchanges every 48hrs.

At day in vitro (DIV) 3.5, cultures were washed three times with a working volume (0.5mL) of Deprivation Media (Neurobasal and 1% Penicillin-Streptomycin) and deprived of trophic support for 12hrs prior to treatment. Cells were then stimulated with 50 ng/mL BDNF (*9*), 200 nM K252a (*52, 53*), or a combination in deprivation media for specified durations (5min to 72hrs). Maintenance Media was re-applied as a positive control (denoted Rescue), while fresh Deprivation Media was applied to the Continued Deprivation negative control.

### 3. Mass cytometry sample collection

Following stimulation, treatment media was replaced with 500 µL of room-temperature Accutase. Cells were monitored for detachment, mechanically dissociated via P1000 trituration into a single-cell suspension, and filtered through a 70 µm strainer into 500 µL of 4% paraformaldehyde (PFA) in PBS. Total processing time from media removal to fixation was maintained under 90s to preserve transient signaling states as best as possible. After a 10 min fixation at room temperature, cells were pelleted (500*g, 5 min), resuspended in 500 µL of Cell Storage Media (CSM; DPBS with 0.5% w/v BSA and 0.2% w/v sodium azide), and aliquoted for quality control and cell counting. Samples were stored at -80°C until analysis.

Each replicate (n) is one culture well; wells came from 2 to 3 independent timed-pregnant litters, except in the 1hr K252A experiment, in which all wells came from a single litter (table S2). To ensure statistical robustness and account for batch effects, treatments were collected with 2-4 biological replicates per time point (table S2), with BDNF-treated samples (5min to 16hr) performed in triplicate and baseline (0hr) controls in quadruplicate.

### 4. Sample barcoding, staining, and mass cytometry

Samples were stained according to established high-dimensional mass cytometry protocols (*54–58*). To minimize inter-sample staining variability and mitigate batch effects, individual samples were uniquely barcoded using a “6-choose-3” combinatorial system of palladium metals (*59, 60*). To account for potential inter-run variations across barcoded batches, a “universal reference” sample – comprising pooled aliquots (11µL, ∼2.44%) of all experimental samples pooled – was included in every barcode set. Barcoding was achieved through reversible permeabilization in 0.02% saponin for 15 min at room temperature with agitation (*61*).

Following barcoding, samples were pooled and blocked with 10% Normal Donkey Serum in CSM for 30min. C were first stained for 30min with a primary antibody cocktail targeting extracellular epitopes, including the TrkB extracellular domain that enabled our assessment of surface TrkB dynamics (table S1). Samples were then fixed and permeabilized with ice-cold 100% methanol for 10min, with vortexing every 2min to prevent cell clumping. Permeabilized cells were subsequently stained for 1hr with a cocktail of antibodies against intracellular targets (table S1). To distinguish single-cell events from debris, samples were incubated overnight at 4°C in 1.6% PFA/DPBS containing 125 µM Iridium (Ir)-intercalator (Standard Biotools, South San Francisco, CA).

Prior to analysis, pooled samples were resuspended in Maxpar Cell Acquisition Solution (CAS) Plus and filtered through a 40µm strainer. Data were acquired on a Helios CyTOF XT system (Standard Biotools) at an acquisition rate below 500 events per second. Samples were processed in four sequential batches of 20 barcoded samples each (table S2). Data were collected using Helios CyTOF.

### 5. Normalization, debarcoding, and single cell gating

Raw FCS files were standardized via bead-based normalization to account for instrument sensitivity fluctuations (*62*). Samples were then debarcoded using a “6-choose-3” combinatorial system to assign events to their respective biological replicates (*60*). To ensure high data quality for the resulting dataset, single-cell events were isolated through a standardized cleanup gating strategy in Cytobank.

Events were first filtered by low barcode separation distance and high Mahalanobis distance to remove poorly assigned cells. Singlets were isolated by comparing event center to event length and width. Residual EQ calibration beads were excluded by gating out high cerium (Ce140) events, while environmental contaminants (Sn120, I127, CS133, Ba138, Pb208) were identified and removed by gating against the Iridium DNA intercalator (Ir191/193). A specific gate on Gd157 was applied to eliminate bleed-through signal from Nd141. The resulting high-fidelity ∼680,000-cell dataset of single-cell events were exported as individual FCS files per sample for downstream analysis (https://community.cytobank.org/cytobank/experiments/121988).

### 6. Batch correction

To account for potential batch effects, data from the “universal reference” sample included in each barcode batch were used to normalize signal intensities across batches using a quantile-based adjustment method (https://github.com/CUHIMSR/CytofBatchAdjust; R-4.5.2) (*63*). This approach effectively minimized technical variance (batch effects) while preserving biological heterogeneity across the longitudinal dataset. Although 72hr samples were collected and processed through the batch correction, they were not included in downstream analyses in the present study due to the possibility of post-treatment secretion of growth factors confounding our interpretation of results.

### 7. Leiden clustering and UMAP visualization

Single-cell events were partitioned into molecularly defined subpopulations using the Leiden community detection algorithm (Python 3.12.4) (*56, 64, 65*). We employed a two-stage clustering strategy to decouple biological identity from signaling state. First, cells were clustered using lineage-specific ID markers (table S1) to define cell-type identities (Fig. 1B, fig. S1). Second, to identify unsupervised signaling signatures (Fig. 4), events – excluding those classified as ‘Unassigned’ – were re-clustered based solely on phosphoprotein markers without a priori identity information. Lineage assignments were subsequently mapped back to signaling clusters using unique event identifiers to determine the cell-type composition of each signature.

High-dimensional data were dimensionally reduced and visualized via Uniform Manifold Approximation and Projection (UMAP) (*66, 67*) using the following parameters: nearest neighbors = 15, metric = Euclidean, local connectivity = 1, epochs = 1000. UMAP coordinates were generated using Python 3.12.4 and visualized with R-4.5.2.

### 8. Data analysis and quantification

Pre-processed mass cytometry data were analyzed using custom R (R-4.5.2) scripts. To quantify signaling induction across the heterogeneous population, we defined a “signaling response” as any event exceeding the 95th percentile of the baseline signal intensity observed in the 0-hour (trophic-deprived) control for each respective phosphoprotein. This thresholding was applied globally and, where indicated, to individual cell-type clusters to account for lineage-specific baseline variances (fig. S2).

Responsiveness was quantified as percent abundance, representing the proportion of “responsive” cells relative to the total population within a given treatment condition or cell-type cluster. For stoichiometry analysis, receptor expression tiers (“None”, “Lo”, “Med”, “Hi”) were established via logical biaxial gating of total TrkB and p75NTR across the entire dataset. This tiered approach allowed for the identification of the “progenitor paradox” and supported the finding that receptor density alone cannot explain signaling output. All custom analysis scripts and data processing pipelines are available at https://github.com/TooSewellForSkool/Sewell-et-al-BDNF-Signaling-Manuscript.

### 9. Statistics

Data are presented as mean ± standard deviation. Replicates (n) are one culture well (sample), reduced to a single value per marker. Groups contained 2-4 samples, derived from 2-3 litters (1 litter for the 1hr K252A experiment), and samples were treated as independent in all analyses; the litter of each sample is listed in table S2.

For time courses, each cluster at each time point was compared with the NoTreat 0hr of the same cluster. For each treatment and marker, Welch’s ANOVA tested for any change across time points, and each time point was then compared with the reference using Welch’s unpaired two-tailed t-test (equal variances is not assumed). P-values were adjusted using the Holm method. For comparisons between conditions at a single time point (Figs 2B, 3F, S4A, S6C, S7B), Welch’s t-tests were applied with Holm adjustment across the three pairs.

Groups with fewer than three samples (n = 2) are plotted without SD bars or significance marks. Their comparisons are reported in tables S3 to S8 for completeness but are underpowered and do not inform conclusions in the main text; because they are included in each Holm family, they make the adjustment of the other time points more conservative. Significance marks are based on the adjusted p-value (* p < 0.05, ** p < 0.01, *** p < 0.001). Unadjusted and adjusted p-values, group means, and n are given in tables S3 to S8. All statistical analyses were performed in R-4.5.2.

## Supplementary Materials

**fig. S1.**
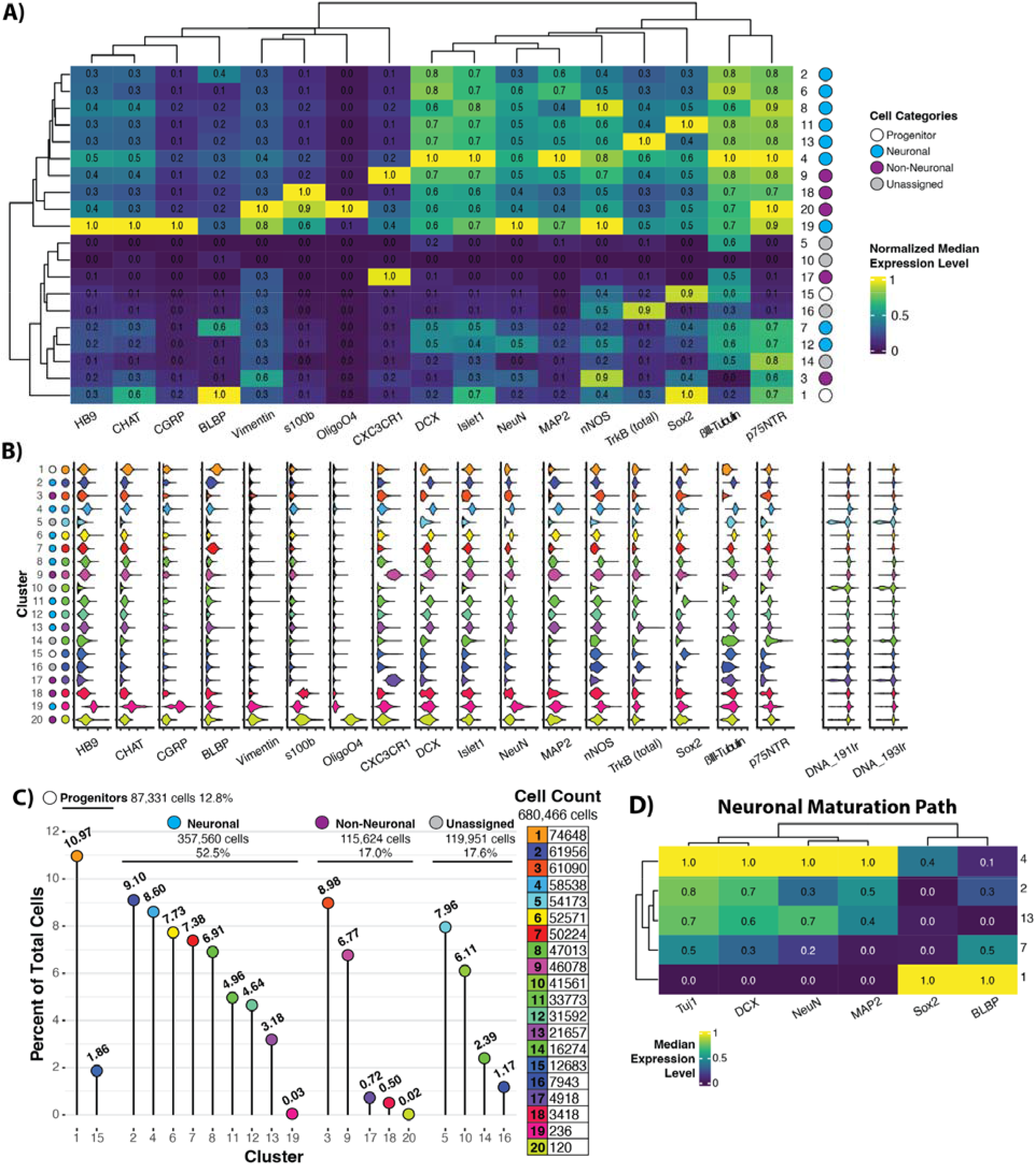
Characterization of cell identity clusters and their distribution. (**A**) Expression heatmap of cell ID markers used for Leiden clustering. (**B**) Violin plots depicting the level of each cell ID marker across each cluster. (**C**) Cell category and cluster distribution across the 680,466 cells measured within the dataset. (**D**) Expression heatmap highlighting the neuronal maturation path that is observed. For expression heatmaps, values represent median expression levels normalized from 0 to 1 for each marker, and dendrograms represent unsupervised grouping of similar markers and clusters.

**fig. S2.**
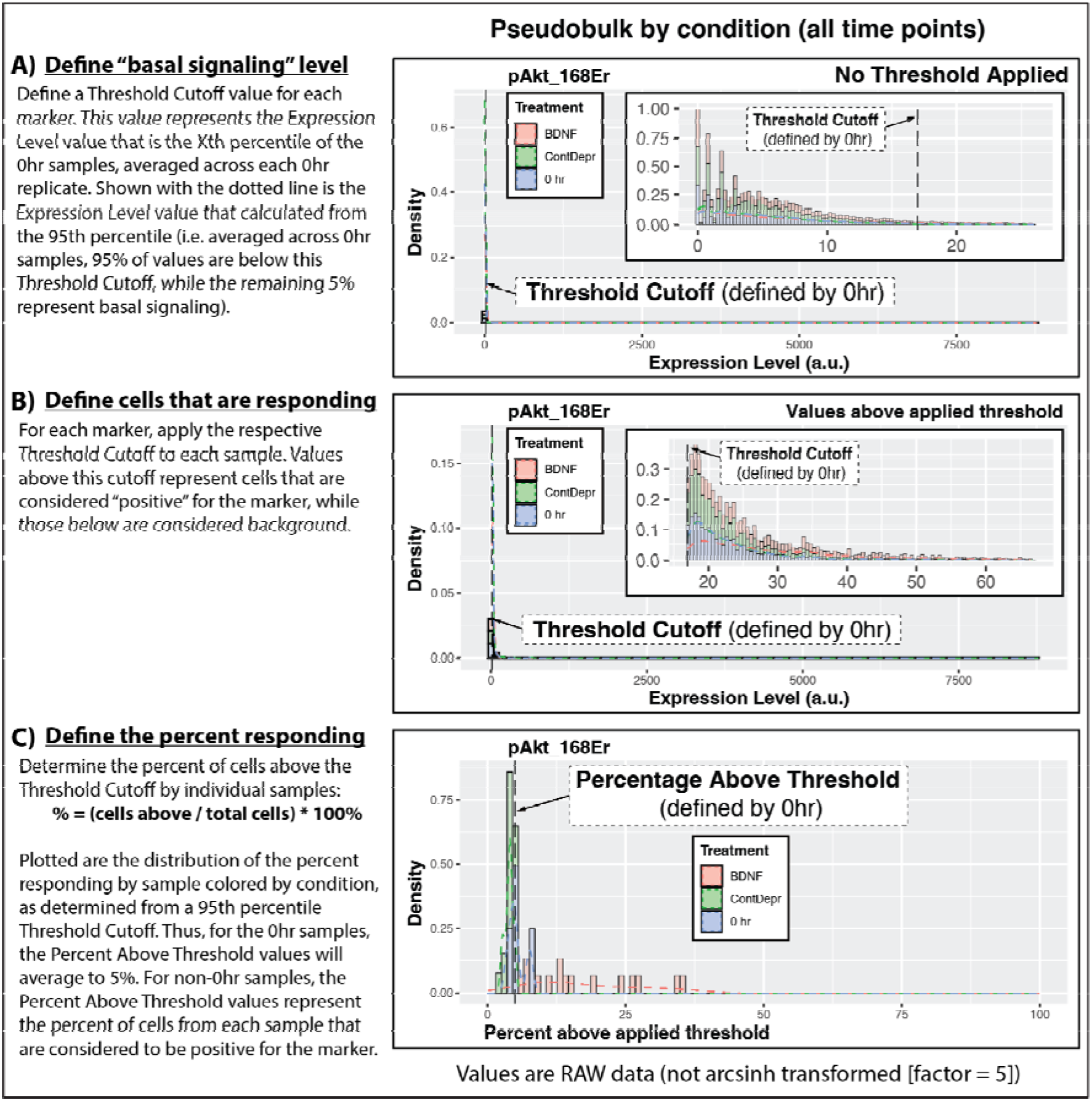
Defining the cells that are responsive to treatment. (**A**) The signaling at time = 0hr represents the baseline (”basal signaling”) to compare against. For each signaling marker, we delineate “real” response from background at time = 0hr based on a threshold cutoff calculated from a defined percentile of the marker abundance. (**B**) Per marker, this threshold cutoff is applied to each sample (across treatment condition, time, and replicate), thus delineating “real” response from background in every measured sample. (C) Per marker, the percent of cells exhibiting “real” response is determined. This percentage is normalized to time = 0hr and reported as the fold change.

**fig. S3.**
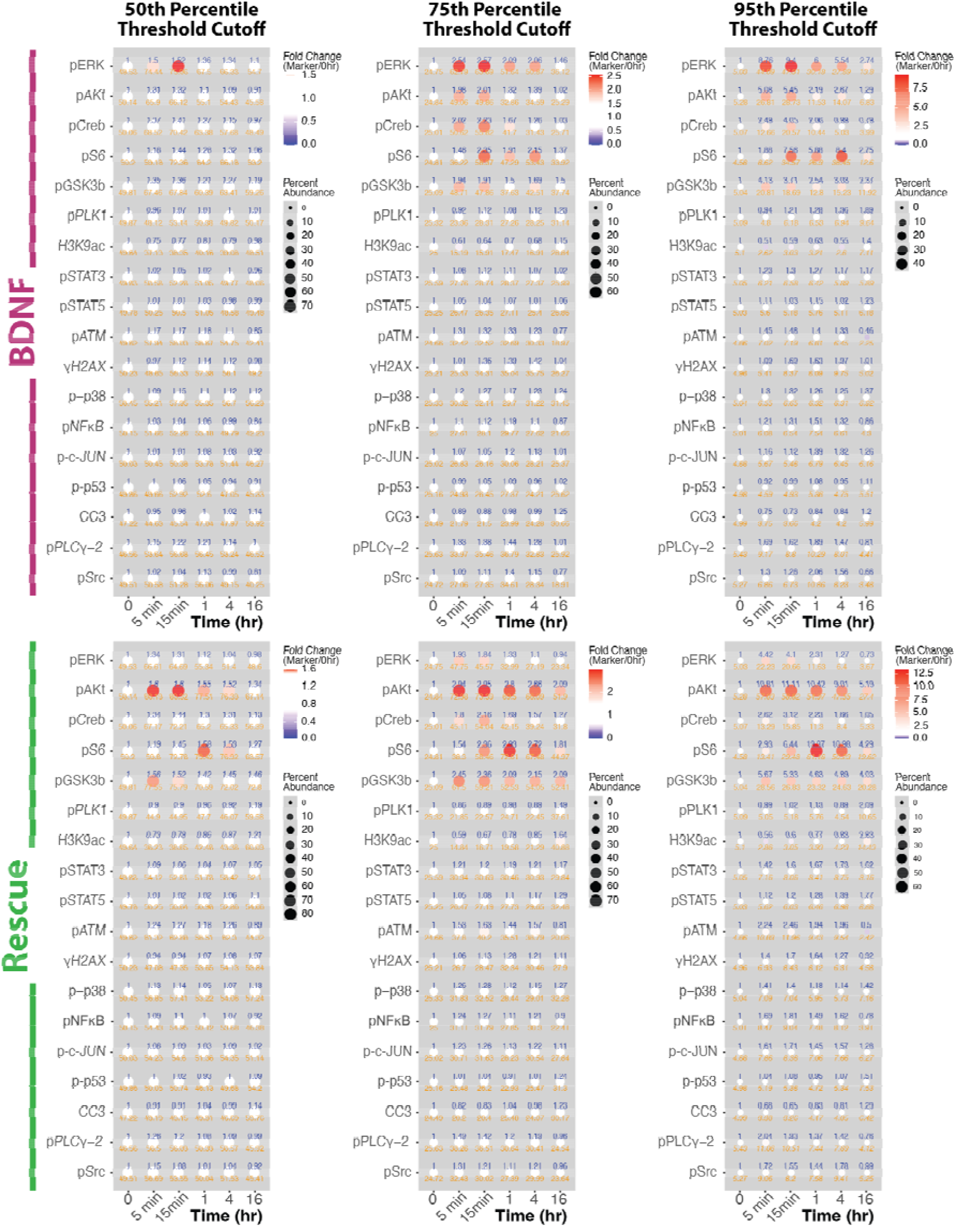
Comparison of different threshold cutoffs and different treatments. Responsive cells defined by a (**A**) 50th, (**B**) 75th, and (**C**) 95th percentile threshold cutoff for BDNF and complete-medium treatment. Dots represent the mean of biological replicates (0hr, n = 4; Rescue, n = 2-3; BDNF, n = 3). Size (orange) indicates the percent of cells in a given sample that contribute the observed response. Color (blue) indicates the fold change relative to time = 0hr.

**fig. S4.**
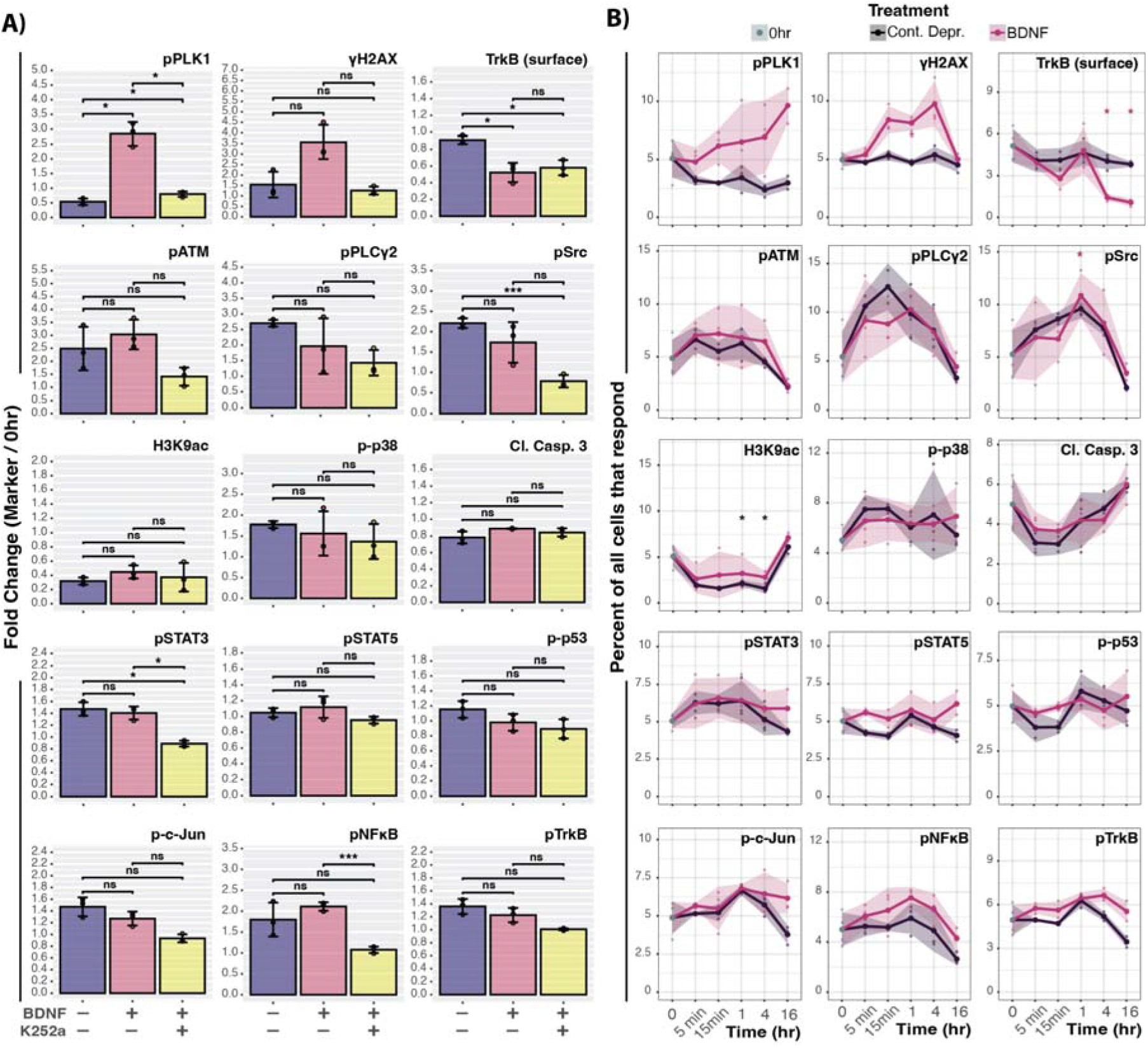
BDNF-induced responses and the effect of K252A inhibition – Extension of Figure 2. (**A**) The fold change relative to 0hr of control (–/–; +BDNF) and co-treatment of BDNF with a Trk inhibitor (K252A) at 1hr (n = 3 per condition). (**B**) Percent abundance plotted for individual signaling markers comparing continued deprivation (Cont. Depr.) and BDNF. Each data point represents the percent of all cells in the collected sample that contributes to the measured response. (A) Points represent mean±SD of 3 culture wells (n = 3) from 1 litter. (B) Points represent mean±SD of 2-4 culture wells (0hr, n = 4; Cont. Depr., n = 2-3; BDNF, n = 3) from 2-3 litters. p-values are adjusted for multiple comparisons (Welch’s unpaired two-sided t-tests with Holm adjustment): * p < 0.05, ** < 0.01, *** < 0.001; ns = not significant.

**fig. S5.**
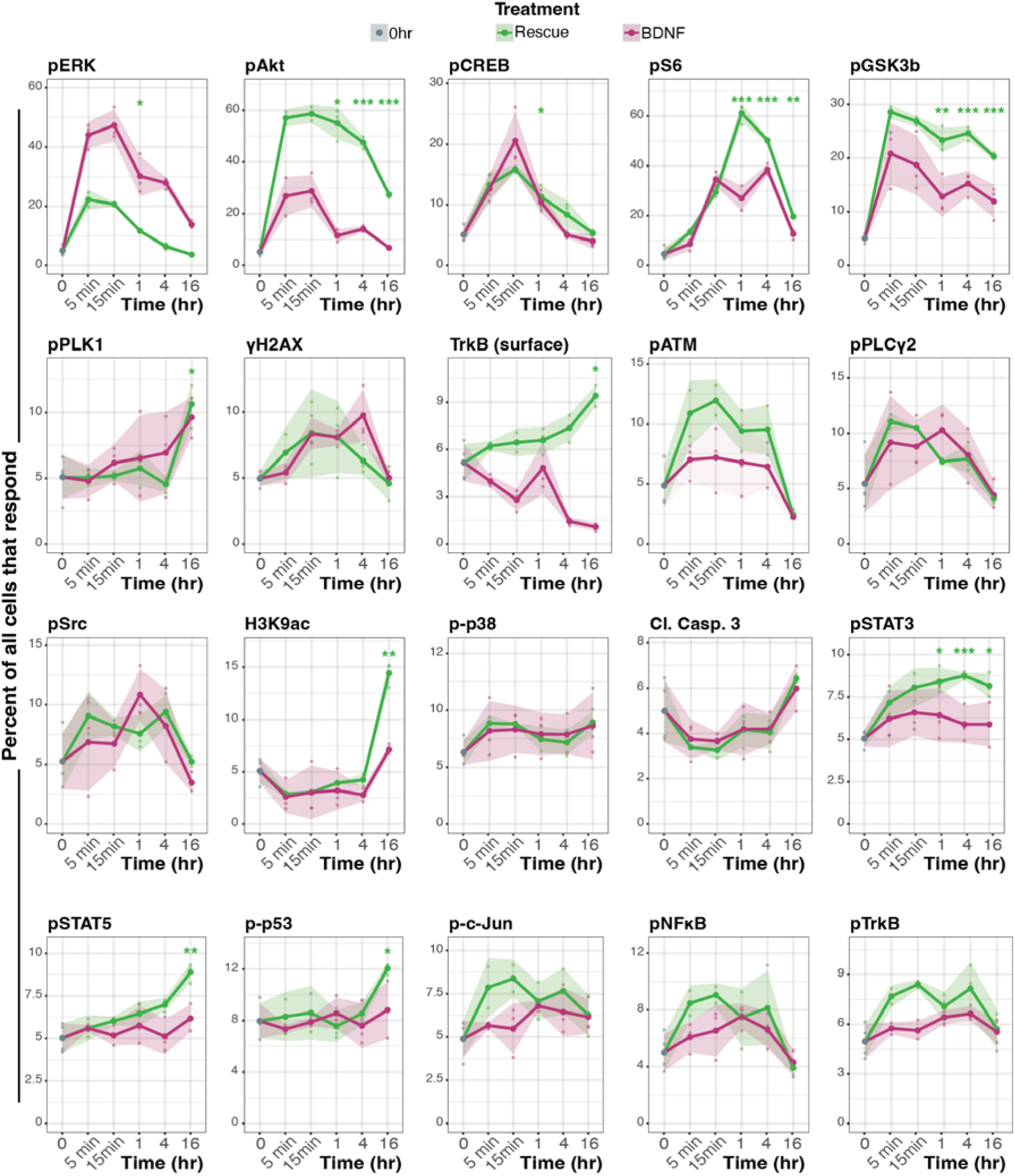
Pseudobulk signaling response to Rescue and BDNF treatment. Line graphs show pseudobulk analysis of the percent of all cells that respond to Rescue or BDNF stimulation across each signaling marker. BDNF data are the same samples shown in Fig. 2C and fig. S4B, replotted here for visual comparison; significance is shown for Rescue only, and statistics for BDNF are given with Fig. 2C (table S3). Rescue and BDNF were not compared statistically. Points represent mean±SD of 2-4 culture wells (0hr, n = 4; Cont. Depr., n = 2-3; BDNF, n = 3) from 2-3 litters. p-values are adjusted for multiple comparisons (Welch’s unpaired two-sided t-tests with Holm adjustment): * p < 0.05, ** < 0.01, *** < 0.001; ns = not significant. See Supplemental Table 3 for values.

**fig. S6.**
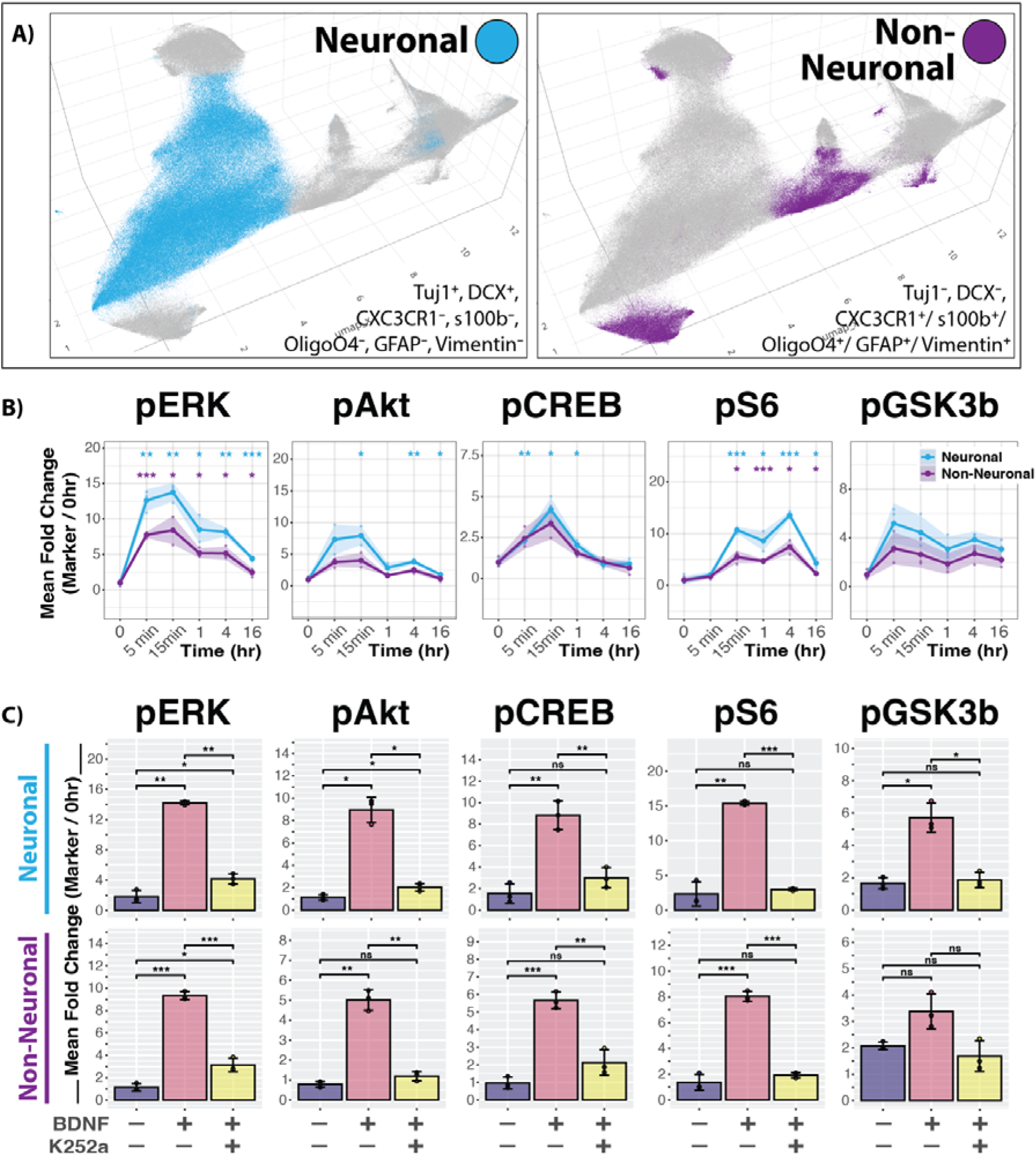
BDNF-induced signaling changes in Neuronal and Non-Neuronal cell categories. (**A**) 3D UMAP visualization of the cells within each category. (**B**) Line graphs compare signaling responses to BDNF across cell categories. (**C**) Bar graphs depict the relative effects of K252A treatment at 1hr. (B) Points represent mean±SD of 2-4 culture wells (n) from 2-3 litters. (C) Points represent mean±SD of 3 culture wells (n = 3) from 1 litter. p-values are adjusted for multiple comparisons (Welch’s unpaired two-sided t-tests with Holm adjustment): * p < 0.05, ** < 0.01, *** < 0.001; ns = not significant. See Supplemental Table 4 for values.

**fig. S7.**
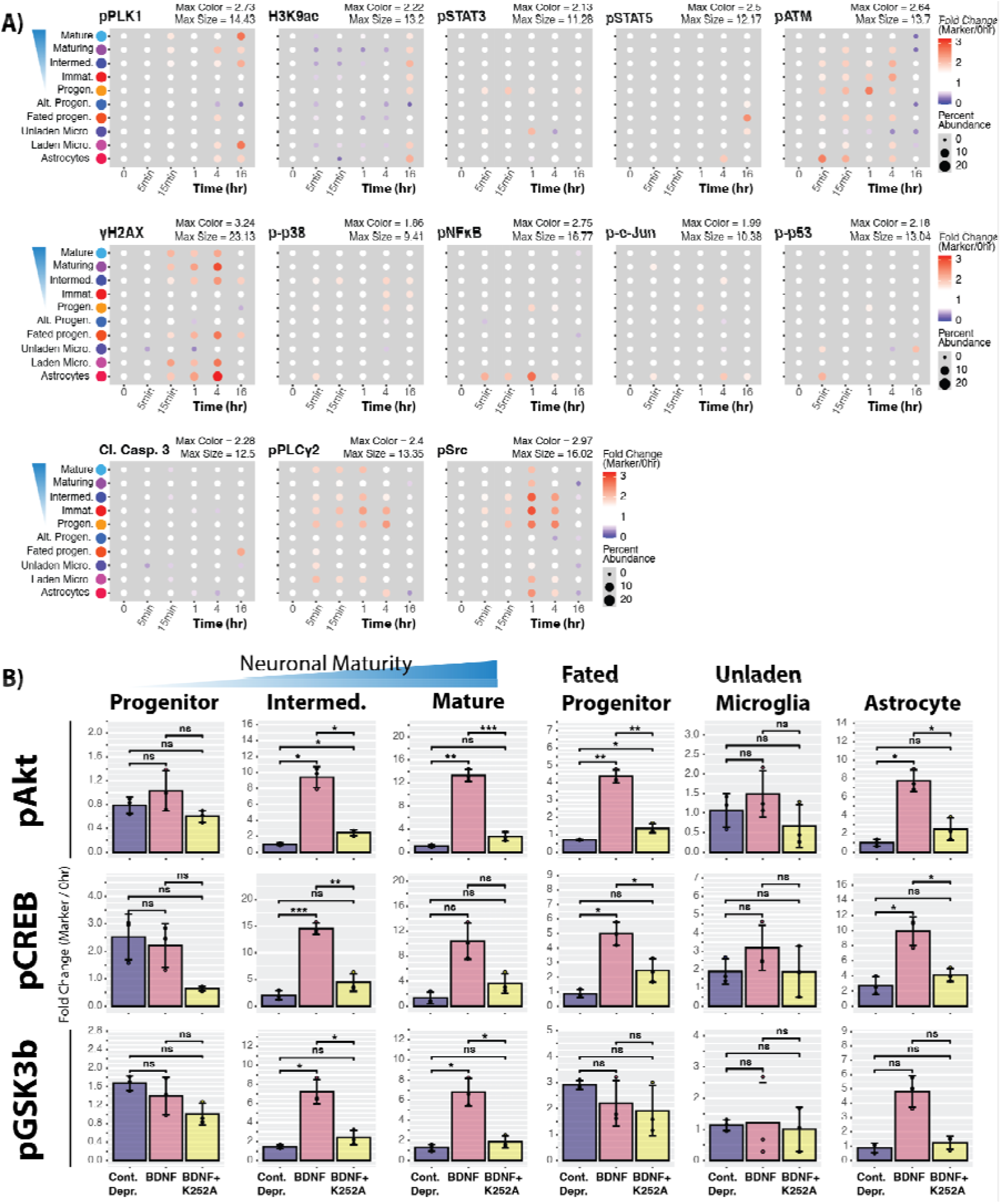
BDNF-induced signaling across distinct cell types – Extension of Fig. 3. (**A**) Dot plots depicting the percent of cells that increase activation of signaling molecules not included in Figure 3 within each identified cell type. Size indicates the percent of the given cell type that contribute to the response, while color indicates the mean fold change relative to 0hr (0hr, n = 4; Rescue, n = 2-3; BDNF, n = 3) (**B**) Bar graphs depict the relative effects of Trk inhibition on BDNF-induced signaling across cell types (expanding on the markers shown in Fig. 3). Bars represent mean±SD of 3 culture wells (n = 3) from 1 litter; p-values are adjusted for multiple comparisons (Welch’s unpaired two-sided t-tests with Holm adjustment): * p < 0.05, ** < 0.01, *** < 0.001; ns = not significant. See Supplemental Table 5 for values.

**fig. S8.**
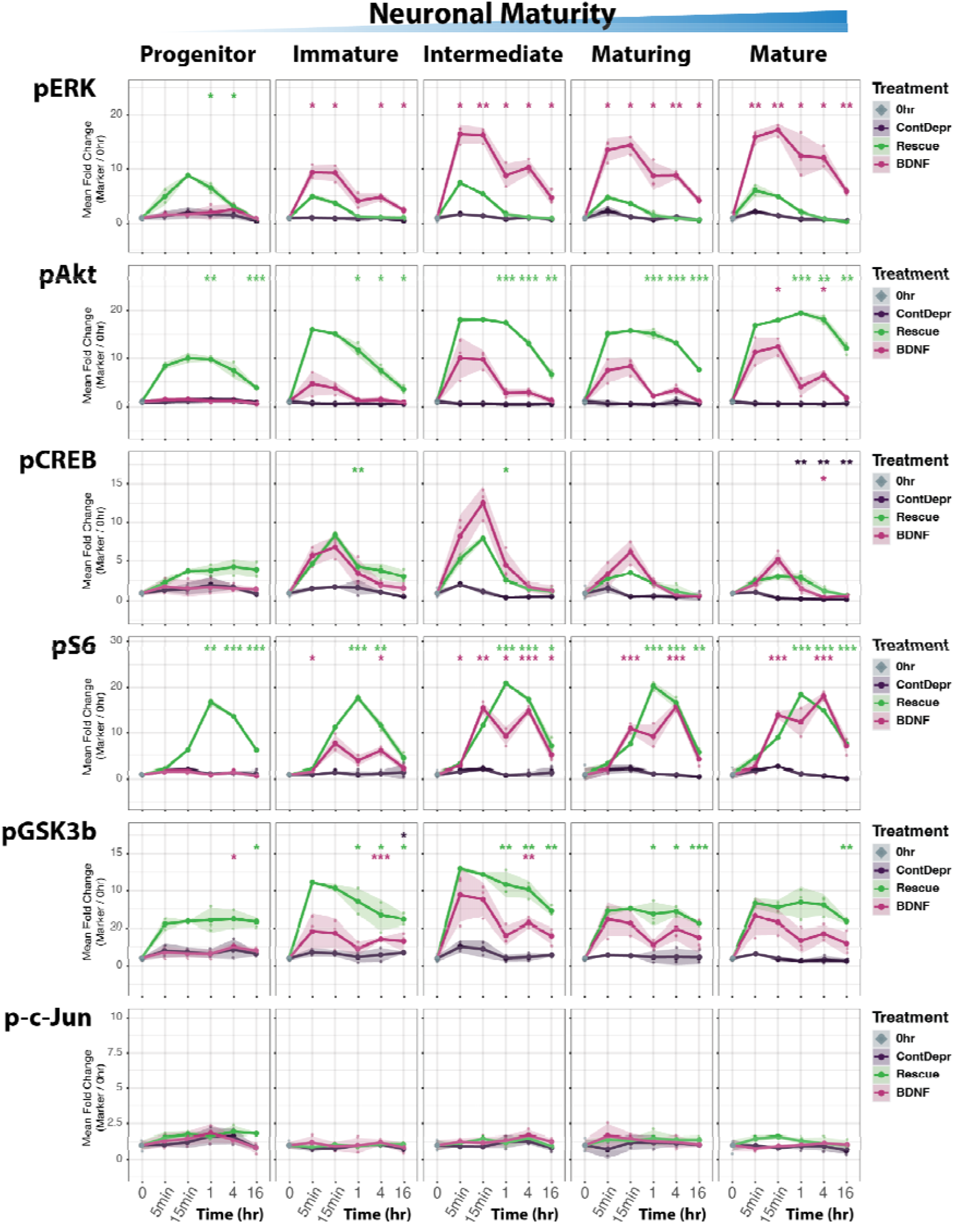
Comparison of signaling responses across treatments in neuronal lineage cell types. Line graph comparison of continued deprivation control, complete-medium rescue, and BDNF treatments of selected markers for clusters identified in the neuronal maturation path. Points represent mean±SD of 2-4 culture wells (0hr, n = 4; Cont. Depr., n = 2-3; BDNF, n = 3) from 2-3 litters. p-values are adjusted for multiple comparisons (Welch’s unpaired two-sided t-tests with Holm adjustment): * p < 0.05, ** < 0.01, *** < 0.001; ns = not significant. See Supplemental Table 6 for values.

**fig. S9.**
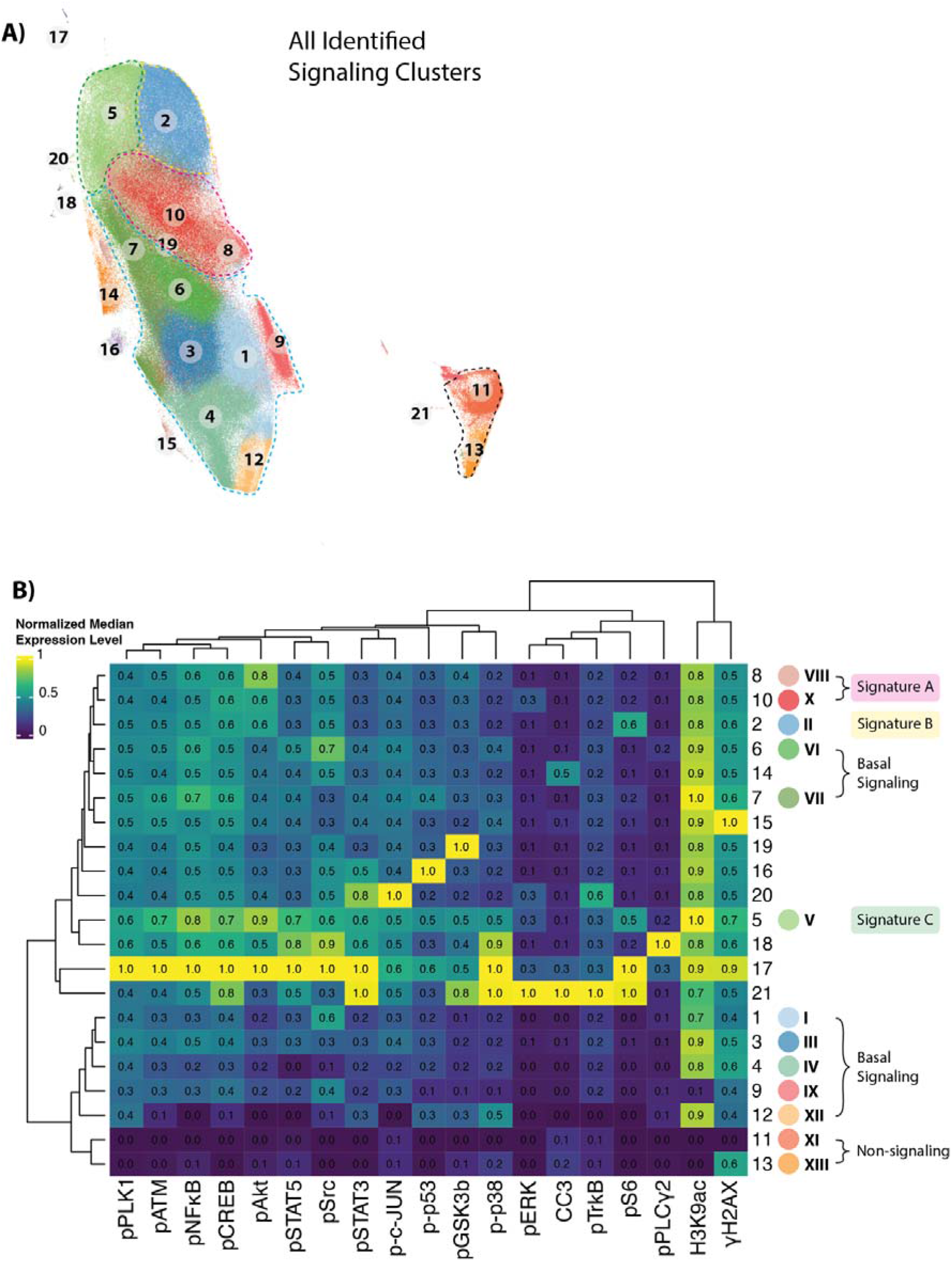
Signaling signatures agnostic to cell ID clustering information. (**A**) UMAP visualization of signaling clusters identified by using only signaling markers for Leiden clustering analysis. (**B**) Expression heatmap molecularly defines signaling signatures identified by clustering analysis.

**fig. S10.**
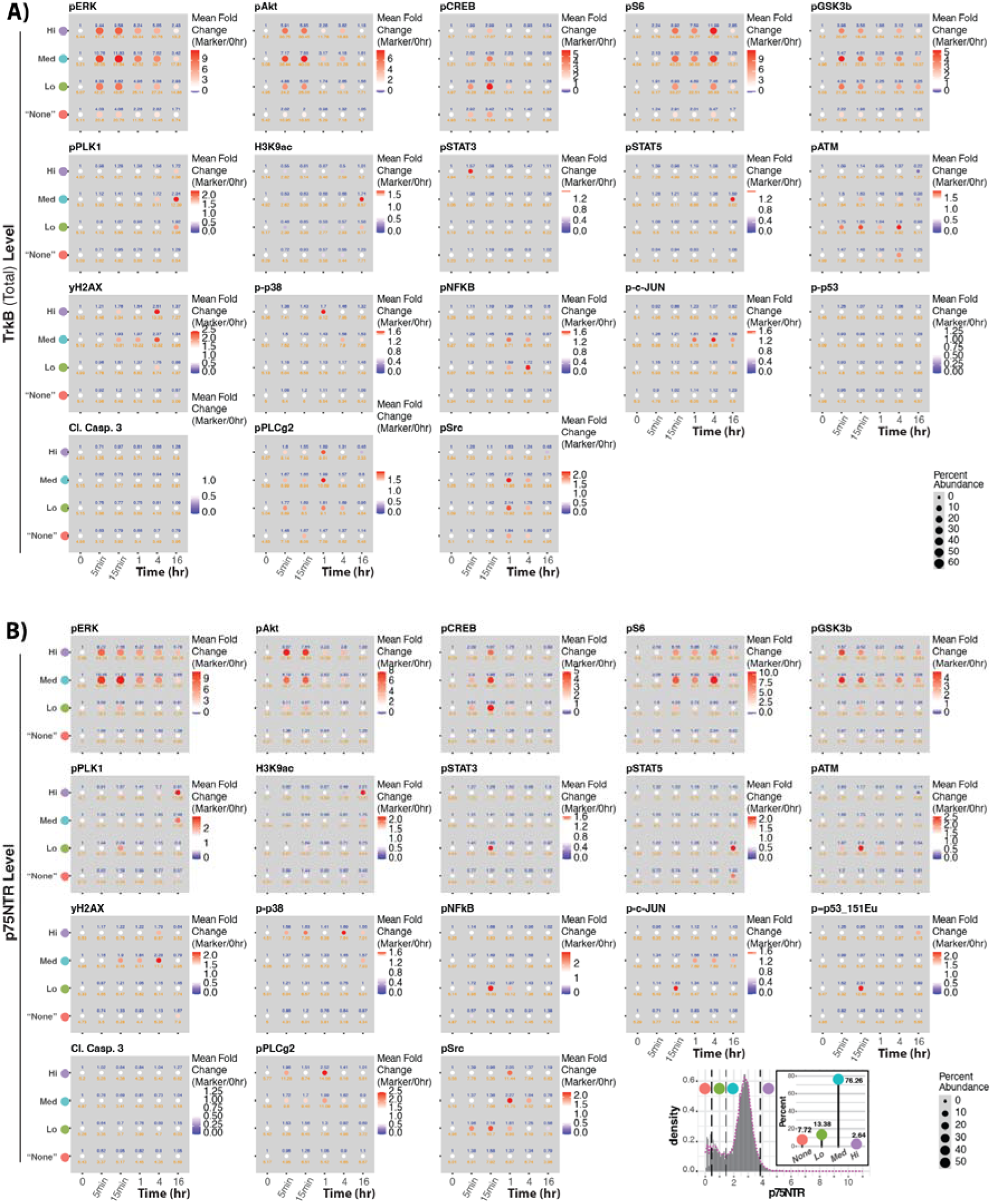
BDNF receptor level contribution to pseudobulk signaling response. Pseudobulk signaling response across signaling markers categorized based on (**A**) TrkB (total) or (**B**) p75NTR level – Distribution and categorization of p75NTR expression level in Pseudobulk data (Inset depicts percent of all cells within each expression category). Quantitative fold change values (color) are denoted in blue. Quantitative percent abundance values (size) are denoted in orange.

**fig. S11.**
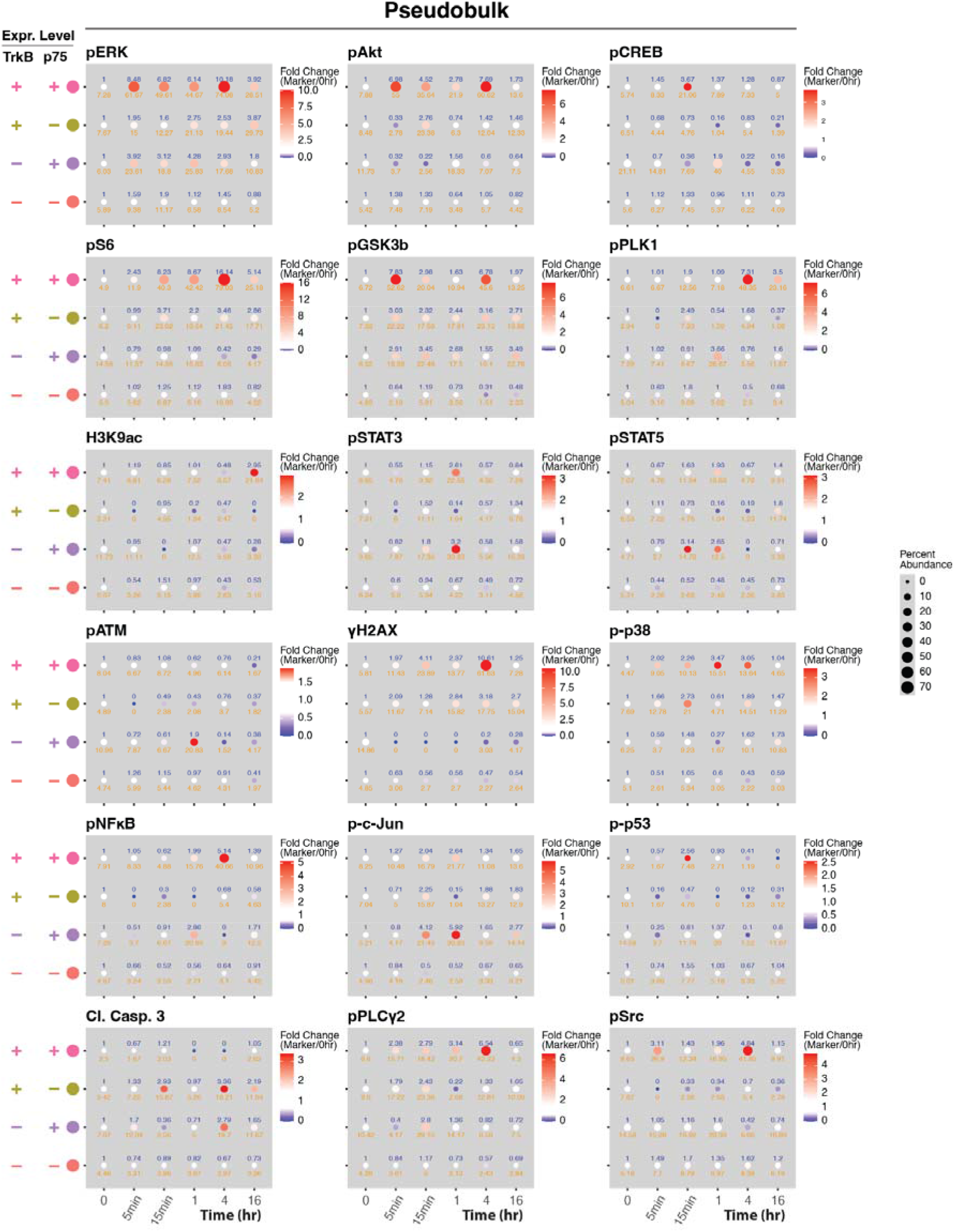
Pseudobulk analysis of receptor level contribution to signaling heterogeneity. Dot plots depict signaling dynamics across TrkB expression comparing p75+/– at the same TrkB expression level. Quantitative fold change values (color) are denoted in blue. Quantitative percent abundance values (size) are denoted in orange.

**fig. S12.**
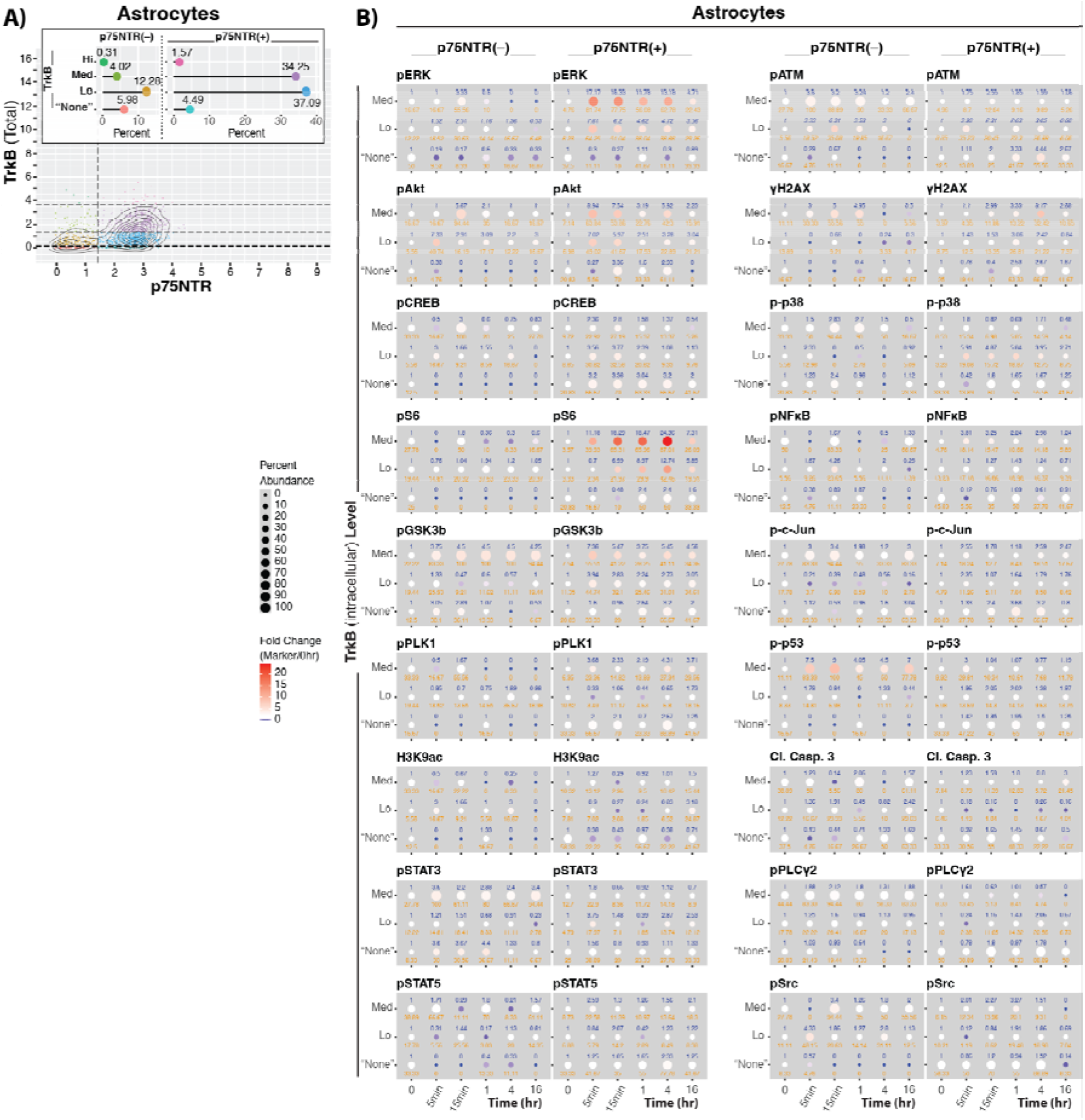
Receptor expression level contributes to signaling heterogeneity within the same cell type. (**A**) Distribution and categorization of Astrocytic cells based on TrkB, p75NTR co-/expression. (**B**) Dot plots depict signaling dynamics across TrkB expression comparing p75+/– at the same TrkB expression level. Quantitative fold change values (color) are denoted in blue. Quantitative percent abundance values (size) are denoted in orange.

**fig. S13.**
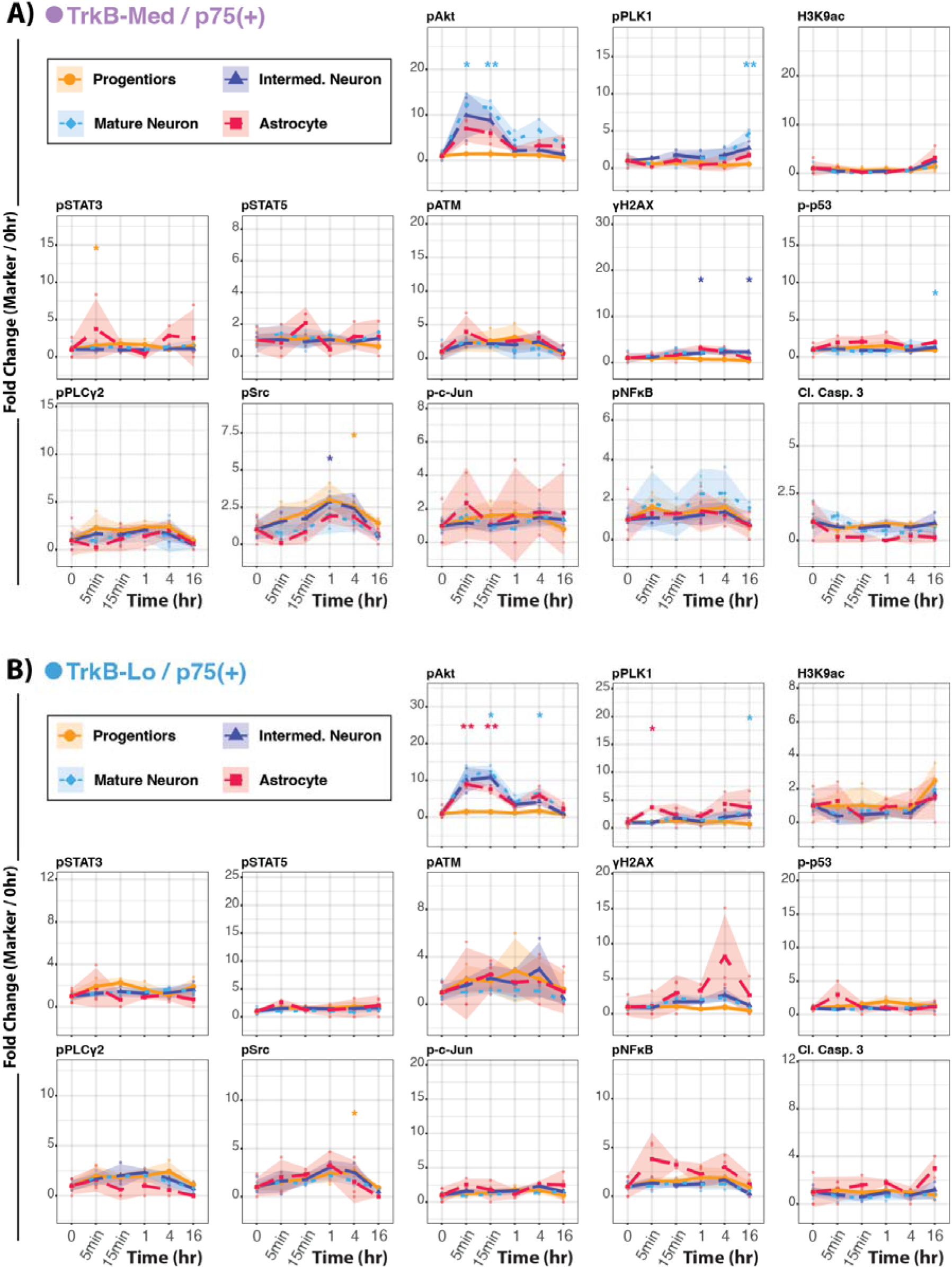
Signaling comparison of different cell types with the same receptor profile – Extension of Fig. 6D, E. Line graphs compare signaling responses in different cell types with the same (**A**) TrkB-Med/p75NTR+ or (**B**) TrkB-Lo/p75NTR+ receptor expression profiles. Points represent mean±SD of 2-4 culture wells (0hr, n = 4; Cont. Depr., n = 2-3; BDNF, n = 3) from 2-3 litters. p-values are adjusted for multiple comparisons (Welch’s unpaired two-sided t-tests with Holm adjustment): * p < 0.05, ** < 0.01, *** < 0.001; ns = not significant. See Supplemental Table 8 for values.

**Table S1. Mass cytometry antibody panel.** Cells were stained with 38 antibodies representing cell ID (18) and cell signaling (19) markers. Additionally, a TrkB antibody was used to stain surface receptors, to measure the reduction of surface TrkB as a correlate of internalization. Antibodies were either purchased in a conjugated form or conjugated in lab using a MaxPar X8 Antibody Labeling kit. Select markers were used for clustering analysis to determine cell ID clusters (Fig. 1) and signaling clusters (Fig. 4). Stock concentrations are in mg/mL and final concentrations in ng/mL. Antibodies purchased as test-sized vials are given in test units: 50x denotes a vial of 50 tests, and 1x or 1/3x the fraction of one test used per sample. * ‘Variable’ indicates the marker was not included in the Leiden clustering analysis: TrkB (surf) excluded because it is variable across time; GFAP excluded because it did not accurately define the correct cell populations.

**Table S2. Sample organization into barcode sets.** The “6-choose-3” combinatorial system of barcoding with palladium metals allows for 20 samples to be barcoded at once. To this end, our 80 samples had to be separated into 4 barcode sets. All barcode sets included a Universal sample – pool of a small aliquot of all 80 samples – which is used for batch correction across sets. Barcode set 1-3 included replicate and triplicate samples of the primary treatment time course. Barcode set 4 includes the 1hr K252A treatment sample set. Stock samples filled gaps in barcode set 3 and 4 to ensure that all sets were treated in the same manner to reduce any barcode set variability. Litter identifies the timed-pregnant litter used for each culture by its internal collection label; each label is a distinct litter: 2.1.1 (collected 2024-11-13), 2.1 (2024-12-09), 2.2 (2024-12-14), 3.3 (2025-02-24) and 3.2.2 (2025-03-17). All samples in barcode set 4 came from litter 3.3. Stock samples were not included in the analyses.

**Table S3. Statistics for pseudobulk analyses (Fig. 2B, C**; **figs. S4A, B, S5).** Each block is labeled with the panel(s) it supports, and Cluster 0 denotes all cells. In the Fig. 2B/S4A block, mean values are the percent of cells above the 95th percentile of the 0hr signal, expressed as fold change relative to 0hr (Scale = FC). The Fig. 2C/S4B and fig. S5 blocks report the percent of cells above threshold (Scale = RAWvalues). Time course blocks (Fig. 2C, figs. S4B, S5): each time point was compared with the NoTreat 0hr samples of the same cluster by Welch’s unpaired two-sided t-test, with Holm adjustment across time points within each treatment and marker (p_primary); Omnibus_p is a Welch ANOVA across time points (blank where it could not be computed, e.g. when a group had zero variance). K252A block (Fig. 2B, fig. S4A): the three 1hr groups (ContDepr, BDNFTest, BDNF+K252aTest) were compared pairwise by Welch’s t-test with Holm adjustment across the three pairs (p_Holm, repeated as p_primary so that every block has the same primary column); Omnibus_p is a Welch ANOVA and Omnibus_p_classic a classic one-way ANOVA across the three groups, and p_Tukey is Tukey’s HSD, given for reference. All K252A samples came from a single litter (table S2). The fig. S5 block covers complete-medium rescue (CompMed) only; BDNF statistics for the same samples are in the Fig. 2C/S4B block. In all tables, p_unadjusted is the unadjusted p-value; q_BH_all_markers is a Benjamini-Hochberg q-value across the 22 signaling markers and all time points (or pairs) within a treatment and cluster, given as an alternative correction, and Family_Size_BH is the number of tests in that family. Significance marks p_primary: * < 0.05, ** < 0.01, *** < 0.001. N_Group and N_Reference are numbers of culture wells. Comparisons involving groups with two culture wells (n = 2) are reported for completeness but are underpowered and do not inform conclusions in the main text; because they are included in each Holm family, they make the adjustment of the other time points more conservative. The Significance column is filled for every row, but no marks are shown in the figures for n = 2 groups.

**Table S4. Statistics for cell categories (fig. S6B, C).** Cluster denotes cell category (1 = progenitor, 2 = neuronal, 3 = non-neuronal, 4 = unassigned). Mean values are the percent of cells above the 95th percentile of the 0hr signal, expressed as fold change relative to 0hr (Scale = FC). Time course block (fig. S6B, BDNF): each time point was compared with the NoTreat 0hr samples of the same cluster by Welch’s unpaired two-sided t-test, with Holm adjustment across time points within each treatment and marker (p_primary); Omnibus_p is a Welch ANOVA across time points (blank where it could not be computed, e.g. when a group had zero variance). K252A block (fig. S6C): the three 1hr groups (ContDepr, BDNFTest, BDNF+K252aTest) were compared pairwise by Welch’s t-test with Holm adjustment across the three pairs (p_Holm, repeated as p_primary so that every block has the same primary column); Omnibus_p is a Welch ANOVA and Omnibus_p_classic a classic one-way ANOVA across the three groups, and p_Tukey is Tukey’s HSD, given for reference. All K252A samples came from a single litter (table S2). Columns are as described for table S3.

**Table S5. Statistics for cell ID clusters (Fig. 3C-F; fig. S7B).** Cluster refers to the cell ID clusters in Fig. 1C and fig. S1. Mean values are the percent of cells above the 95th percentile of the 0hr signal, expressed as fold change relative to 0hr (Scale = FC); a value of 0 indicates that no cells exceeded the threshold in any replicate. Time course blocks (Fig. 3C, E: BDNF; Fig. 3D: complete-medium rescue, CompMed): each time point was compared with the NoTreat 0hr samples of the same cluster by Welch’s unpaired two-sided t-test, with Holm adjustment across time points within each treatment and marker (p_primary); Omnibus_p is a Welch ANOVA across time points (blank where it could not be computed, e.g. when a group had zero variance). K252A block (Fig. 3F, fig. S7B): the three 1hr groups (ContDepr, BDNFTest, BDNF+K252aTest) were compared pairwise by Welch’s t-test with Holm adjustment across the three pairs (p_Holm, repeated as p_primary so that every block has the same primary column); Omnibus_p is a Welch ANOVA and Omnibus_p_classic a classic one-way ANOVA across the three groups, and p_Tukey is Tukey’s HSD, given for reference. All K252A samples came from a single litter (table S2). Columns are as described for table S3.

**Table S6. Statistics for treatment comparisons in neuronal lineage cell types (fig. S8).** One block per cell ID cluster (Fig. 1C; fig. S1), each covering continued deprivation (ContDepr), complete-medium rescue (CompMed) and BDNF. Mean values are the percent of cells above the 95th percentile of the 0hr signal, expressed as fold change relative to 0hr (Scale = FC). For each treatment, each time point was compared with the NoTreat 0hr samples of the same cluster by Welch’s unpaired two-sided t-test, with Holm adjustment across time points within each treatment and marker (p_primary); Omnibus_p is a Welch ANOVA across time points (blank where it could not be computed, e.g. when a group had zero variance). BDNF rows repeat those in table S5 (Fig. 3C, E block), as do CompMed rows for cluster 1 (Fig. 3D block). Columns are as described for table S3.

**Table S7. Statistics for surface TrkB (Fig. 5F).** One block per treatment (ContDepr, BDNF, CompMed), giving surface TrkB (TrkB_surface) in each cell ID cluster (Fig. 1C; fig. S1). Values are the mean surface TrkB signal relative to 0hr (Scale = FC), not the percent of cells above threshold used in the other tables. For each treatment, each time point was compared with the NoTreat 0hr samples of the same cluster by Welch’s unpaired two-sided t-test, with Holm adjustment across time points within each treatment and marker (p_primary); Omnibus_p is a Welch ANOVA across time points (blank where it could not be computed, e.g. when a group had zero variance). Clusters 19 and 20 contained fewer than 15 cells at 0hr; p-values are blank where a group had a single well. Columns are as described for table S3.

**Table S8. Statistics for cell types sharing a receptor profile (Fig. 6D, E; fig. S13).** One block per receptor category (ThreshCat): 23, TrkB-Med/p75+ (Fig. 6D, fig. S13A), and 22, TrkB-Lo/p75+ (Fig. 6E, fig. S13B). Cluster refers to the cell ID clusters in Fig. 1C and fig. S1. Mean values are the percent of cells above the 95th percentile of the 0hr signal, expressed as fold change relative to 0hr (Scale = FC). For BDNF, each time point was compared with the NoTreat 0hr samples of the same cluster by Welch’s unpaired two-sided t-test, with Holm adjustment across time points within each treatment and marker (p_primary); Omnibus_p is a Welch ANOVA across time points (blank where it could not be computed, e.g. when a group had zero variance). Columns are as described for table S3.

## Acknowledgments

We would like to thank all members of the Deppmann, Winckler, and Zunder labs for their helpful discussions, and Dr. Alban Gaultier for recognizing the potential of this project. We thank Isaiah Swann of the Winckler lab for assistance in antibody conjugation. We also thank The University of Virginia Gilmer Hall vivarium staff for assistance with animal care and procurement. We want to acknowledge that the data for this manuscript were generated in the University of Virginia Flow Cytometry Core Facility (RRid:SCR_017829) and is partially supported by the NCI Grant (P30-CA044579), and we thank the core for their expertise and continued support. We also acknowledge Research Computing at The University of Virginia for providing computational resources and technical support that have contributed to the results reported within this manuscript. Figures were partly created in BioRender under a license granted to JMS (https://BioRender.com/5hgxcuy). This was supported by the NIH (JMS, F31NS135970 and T32GM139787; ERZ, 5R01NS111220; BW, R01NS083378; CDD, 5R01NS136342) and the UVA Brain Institute (JMS, Presidential Fellowship in Collaborative Neuroscience). Lastly, we acknowledge the use of AI-assisted technologies in the writing process: JMS used Microsoft Copilot to edit R scripts for clarity and conciseness and Google Gemini to edit the manuscript text for clarity and structure; CDD used Anthropic’s Claude to provide additional edits to the manuscript text, focusing on journal fit; Authors reviewed and edited each component of the manuscript after applying these tools and fully accept responsibility for the contents of this work.

## Author contributions

Conceptualization: JMS, ERZ, BW, CDD

Data Curation: JMS

Formal Analysis: JMS, ACB, GL

Funding Acquisition: JMS, BW, CDD

Investigation: JMS, BW, CDD

Methodology: JMS, ACB

Project Administration: JMS

Resources: JMS, ERZ, BW, CDD

Software: JMS, ERZ

Supervision: ERZ, BW, CDD

Validation: JMS

Visualization: JMS, ACB, GL

Writing – original draft: JMS

Writing – review & editing: JMS, ACB, GL, ERZ, BW, CDD

## Data and materials availability

Raw FCS files are available at https://community.cytobank.org/cytobank/experiments/121988. All custom analysis scripts and data processing pipelines are available at https://github.com/TooSewellForSkool/Sewell-et-al-BDNF-Signaling-Manuscript.

## References and Notes

1. P. Kowiański, G. Lietzau, E. Czuba, M. Waśkow, A. Steliga, J. Moryś, BDNF: A Key Factor with Multipotent Impact on Brain Signaling and Synaptic Plasticity. Cell. Mol. Neurobiol. 38, 579–593 (2018).

2. E. J. Huang, L. F. Reichardt, Neurotrophins: roles in neuronal development and function. Annu. Rev. Neurosci. 24, 677–736 (2001).

3. C. Miranda-Lourenço, L. Ribeiro-Rodrigues, J. Fonseca-Gomes, S. R. Tanqueiro, R. F. Belo, C. B. Ferreira, N. Rei, M. Ferreira-Manso, C. de Almeida-Borlido, T. Costa-Coelho, C. F. Freitas, S. Zavalko, F. M. Mouro, A. M. Sebastião, S. Xapelli, T. M. Rodrigues, M. J. Diógenes, Challenges of BDNF-based therapies: From common to rare diseases. Pharmacol. Res. 162, 105281 (2020).

4. C. S. Wang, E. T. Kavalali, L. M. Monteggia, BDNF signaling in context: From synaptic regulation to psychiatric disorders. Cell. 185, 62–76 (2022).

5. C. Hernández-Del Caño, N. Varela-Andrés, A. Cebrián-León, R. Deogracias, Neurotrophins and their receptors: bdnf’s role in gabaergic neurodevelopment and disease. Int. J. Mol. Sci. 25 (2024), doi:10.3390/ijms25158312.

6. J. Zheng, W.-H. Shen, T.-J. Lu, Y. Zhou, Q. Chen, Z. Wang, T. Xiang, Y.-C. Zhu, C. Zhang, S. Duan, Z.-Q. Xiong, Clathrin-dependent endocytosis is required for TrkB-dependent Akt-mediated neuronal protection and dendritic growth. J. Biol. Chem. 283, 13280–13288 (2008).

7. M. Bibel, E. Hoppe, Y. A. Barde, Biochemical and functional interactions between the neurotrophin receptors trk and p75NTR. EMBO J. 18, 616–622 (1999).

8. R. B. Meeker, K. S. Williams, The p75 neurotrophin receptor: at the crossroad of neural repair and death. Neural Regen. Res. 10, 721–725 (2015).

9. J. P. Zanin, L. E. Montroull, M. Volosin, W. J. Friedman, The p75 neurotrophin receptor facilitates trkb signaling and function in rat hippocampal neurons. Front. Cell. Neurosci. 13, 485 (2019).

10. J. N. Conroy, E. J. Coulson, High-affinity TrkA and p75 neurotrophin receptor complexes: A twisted affair. J. Biol. Chem. 298, 101568 (2022).

11. J. P. S. Makkerh, C. Ceni, D. S. Auld, F. Vaillancourt, G. Dorval, P. A. Barker, p75 neurotrophin receptor reduces ligand-induced Trk receptor ubiquitination and delays Trk receptor internalization and degradation. EMBO Rep. 6, 936–941 (2005).

12. C. Loos, K. Moeller, F. Fröhlich, T. Hucho, J. Hasenauer, A Hierarchical, Data-Driven Approach to Modeling Single-Cell Populations Predicts Latent Causes of Cell-To-Cell Variability. Cell Syst. 6, 593–603.e13 (2018).

13. M. R. Birtwistle, J. Rauch, A. Kiyatkin, E. Aksamitiene, M. Dobrzyński, J. B. Hoek, W. Kolch, B. A. Ogunnaike, B. N. Kholodenko, Emergence of bimodal cell population responses from the interplay between analog single-cell signaling and protein expression noise. BMC Syst. Biol. 6, 109 (2012).

14. M. H. Spitzer, G. P. Nolan, Mass cytometry: single cells, many features. Cell. 165, 780– 791 (2016).

15. M. B. Elowitz, A. J. Levine, E. D. Siggia, P. S. Swain, Stochastic gene expression in a single cell. Science. 297, 1183–1186 (2002).

16. G. Balázsi, A. van Oudenaarden, J. J. Collins, Cellular decision making and biological noise: from microbes to mammals. Cell. 144, 910–925 (2011).

17. J. Hasenauer, C. Hasenauer, T. Hucho, F. J. Theis, ODE constrained mixture modelling: a method for unraveling subpopulation structures and dynamics. PLoS Comput. Biol. 10, e1003686 (2014).

18. T. Andreska, P. Lüningschrör, M. Sendtner, Regulation of TrkB cell surface expression—a mechanism for modulation of neuronal responsiveness to brain-derived neurotrophic factor. Cell Tissue Res. 382, 5–14 (2020).

19. G. Moya-Alvarado, R. Tiburcio-Felix, M. R. Ibáñez, A. A. Aguirre-Soto, M. V. Guerra, C. Wu, W. C. Mobley, E. Perlson, F. C. Bronfman, BDNF/TrkB signaling endosomes in axons coordinate CREB/mTOR activation and protein synthesis in the cell body to induce dendritic growth in cortical neurons. eLife. 12 (2023), doi:10.7554/eLife.77455.

20. L. Salavessa, T. Lagache, V. Malardé, A. Grassart, J.-C. Olivo-Marin, A. Canette, M. Trichet, P. J. Sansonetti, N. Sauvonnet, Cytokine receptor cluster size impacts its endocytosis and signaling. Proc Natl Acad Sci USA. 118 (2021), doi:10.1073/pnas.2024893118.

21. S. G. Dorsey, C. L. Renn, L. Carim-Todd, C. A. Barrick, L. Bambrick, B. K. Krueger, C. W. Ward, L. Tessarollo, In vivo restoration of physiological levels of truncated TrkB.T1 receptor rescues neuronal cell death in a trisomic mouse model. Neuron. 51, 21–28 (2006).

22. L. Tessarollo, S. Yanpallewar, Trkb truncated isoform receptors as transducers and determinants of BDNF functions. Front. Neurosci. 16, 847572 (2022).

23. M. V. Chao, B. L. Hempstead, p75 and Trk: a two-receptor system. Trends Neurosci. 18, 321–326 (1995).

24. A. Vicario, L. Kisiswa, J. Y. Tann, C. E. Kelly, C. F. Ibáñez, Neuron-type-specific signaling by the p75NTR death receptor is regulated by differential proteolytic cleavage. J. Cell Sci. 128, 1507–1517 (2015).

25. Y. Zhang, D. B. Moheban, B. R. Conway, A. Bhattacharyya, R. A. Segal, Cell surface Trk receptors mediate NGF-induced survival while internalized receptors regulate NGF-induced differentiation. J. Neurosci. 20, 5671–5678 (2000).

26. J. L. Urdiales, E. Becker, M. Andrieu, A. Thomas, J. Jullien, L. A. van Grunsven, S. Menut, G. I. Evan, D. Martín-Zanca, B. B. Rudkin, Cell cycle phase-specific surface expression of nerve growth factor receptors TrkA and p75(NTR). J. Neurosci. 18, 6767– 6775 (1998).

27. J. Hsieh, K. Nakashima, T. Kuwabara, E. Mejia, F. H. Gage, Histone deacetylase inhibition-mediated neuronal differentiation of multipotent adult neural progenitor cells. Proc Natl Acad Sci USA. 101, 16659–16664 (2004).

28. D. K. Ma, M. C. Marchetto, J. U. Guo, G. Ming, F. H. Gage, H. Song, Epigenetic choreographers of neurogenesis in the adult mammalian brain. Nat. Neurosci. 13, 1338– 1344 (2010).

29. I. Koppel, T. Timmusk, Differential regulation of Bdnf expression in cortical neurons by class-selective histone deacetylase inhibitors. Neuropharmacology. 75, 106–115 (2013).

30. M. J. Berridge, The AM and FM of calcium signalling. Nature. 386, 759–760 (1997).

31. O. Forostyak, S. Forostyak, S. Kortus, E. Sykova, A. Verkhratsky, G. Dayanithi, Physiology of Ca(2+) signalling in stem cells of different origins and differentiation stages. Cell Calcium. 59, 57–66 (2016).

32. M. D. Bootman, G. Bultynck, Fundamentals of cellular calcium signaling: A primer. Cold Spring Harb. Perspect. Biol. 12 (2020), doi:10.1101/cshperspect.a038802.

33. A. Saghatelyan, Calcium signaling as an integrator and decoder of niche factors to control somatic stem cell quiescence and activation. FEBS J. 290, 677–683 (2023).

34. A. Hoffmann, A. Levchenko, M. L. Scott, D. Baltimore, The IkappaB-NF-kappaB signaling module: temporal control and selective gene activation. Science. 298, 1241–1245 (2002).

35. M. Junkin, A. J. Kaestli, Z. Cheng, C. Jordi, C. Albayrak, A. Hoffmann, S. Tay, High-Content Quantification of Single-Cell Immune Dynamics. Cell Rep. 15, 411–422 (2016).

36. E. R. Bozich, X. Guo, J. L. Wilson, A. Hoffmann, A computational workflow for assessing drug effects on temporal signaling dynamics reveals robustness in stimulus-specific NFκB signaling. PLoS Comput. Biol. 21, e1013344 (2025).

37. R. Guo, D. Han, X. Song, Y. Gao, Z. Li, X. Li, Z. Yang, Z. Xu, Context-dependent regulation of Notch signaling in glial development and tumorigenesis. Sci. Adv. 9, eadi2167 (2023).

38. R. R. Madsen, B. Vanhaesebroeck, Cracking the context-specific PI3K signaling code. Sci. Signal. 13 (2020), doi:10.1126/scisignal.aay2940.

39. L. Goentoro, M. W. Kirschner, Evidence that fold-change, and not absolute level, of beta-catenin dictates Wnt signaling. Mol. Cell. 36, 872–884 (2009).

40. C. S. Cselenyi, E. Lee, Context-dependent activation or inhibition of Wnt-beta-catenin signaling by Kremen. Sci. Signal. 1, pe10 (2008).

41. L. M. Holt, R. D. Hernandez, N. L. Pacheco, B. Torres Ceja, M. Hossain, M. L. Olsen, Astrocyte morphogenesis is dependent on BDNF signaling via astrocytic TrkB.T1. eLife. 8 (2019), doi:10.7554/eLife.44667.

42. D. Puehringer, N. Orel, P. Lüningschrör, N. Subramanian, T. Herrmann, M. V. Chao, M. Sendtner, EGF transactivation of Trk receptors regulates the migration of newborn cortical neurons. Nat. Neurosci. 16, 407–415 (2013).

43. K. J. Martin, N. Shpiro, R. Traynor, M. Elliott, J. S. C. Arthur, Comparison of the specificity of Trk inhibitors in recombinant and neuronal assays. Neuropharmacology. 61, 148–155 (2011).

44. H. Kase, K. Iwahashi, S. Nakanishi, Y. Matsuda, K. Yamada, M. Takahashi, C. Murakata, A. Sato, M. Kaneko, K-252 compounds, novel and potent inhibitors of protein kinase C and cyclic nucleotide-dependent protein kinases. Biochem. Biophys. Res. Commun. 142, 436–440 (1987).

45. M. Cazorla, J. Prémont, A. Mann, N. Girard, C. Kellendonk, D. Rognan, Identification of a low-molecular weight TrkB antagonist with anxiolytic and antidepressant activity in mice. J. Clin. Invest. 121, 1846–1857 (2011).

46. H. K. Teng, K. K. Teng, R. Lee, S. Wright, S. Tevar, R. D. Almeida, P. Kermani, R. Torkin, Z.-Y. Chen, F. S. Lee, R. T. Kraemer, A. Nykjaer, B. L. Hempstead, ProBDNF induces neuronal apoptosis via activation of a receptor complex of p75NTR and sortilin. J. Neurosci. 25, 5455–5463 (2005).

47. N. H. Woo, H. K. Teng, C.-J. Siao, C. Chiaruttini, P. T. Pang, T. A. Milner, B. L. Hempstead, B. Lu, Activation of p75NTR by proBDNF facilitates hippocampal long-term depression. Nat. Neurosci. 8, 1069–1077 (2005).

48. F. Yang, H.-S. Je, Y. Ji, G. Nagappan, B. Hempstead, B. Lu, Pro-BDNF-induced synaptic depression and retraction at developing neuromuscular synapses. J. Cell Biol. 185, 727–741 (2009).

49. A controlled trial of recombinant methionyl human BDNF in ALS: The BDNF Study Group (Phase III). Neurology. 52, 1427–1433 (1999).

50. A. H. Nagahara, M. H. Tuszynski, Potential therapeutic uses of BDNF in neurological and psychiatric disorders. Nat. Rev. Drug Discov. 10, 209–219 (2011).

51. F. A. Schroeder, M. C. Lewis, D. M. Fass, F. F. Wagner, Y.-L. Zhang, K. M. Hennig, J. Gale, W.-N. Zhao, S. Reis, D. D. Barker, E. Berry-Scott, S. W. Kim, E. L. Clore, J. M. Hooker, E. B. Holson, S. J. Haggarty, T. L. Petryshen, A selective HDAC 1/2 inhibitor modulates chromatin and gene expression in brain and alters mouse behavior in two mood-related tests. PLoS ONE. 8, e71323 (2013).

52. P. Perez-Pinera, T. Hernandez, O. García-Suárez, F. de Carlos, A. Germana, M. Del Valle, A. Astudillo, J. A. Vega, The Trk tyrosine kinase inhibitor K252a regulates growth of lung adenocarcinomas. Mol. Cell. Biochem. 295, 19–26 (2007).

53. P. Tapley, F. Lamballe, M. Barbacid, K252a is a selective inhibitor of the tyrosine protein kinase activity of the trk family of oncogenes and neurotrophin receptors. Oncogene. 7, 371–381 (1992).

54. A. B. Keeler, A. L. Van Deusen, I. C. Gadani, C. M. Williams, S. M. Goggin, A. K. Hirt, S. A. Vradenburgh, K. I. Fread, E. A. Puleo, L. Jin, O. Y. Calhan, C. D. Deppmann, E. R. Zunder, A developmental atlas of somatosensory diversification and maturation in the dorsal root ganglia by single-cell mass cytometry. Nat. Neurosci. 25, 1543–1558 (2022).

55. S. A. Vradenburgh, A. L. Van Deusen, A. N. Beachum, J. M. Moats, A. K. Hirt, C. D. Deppmann, A. B. Keeler, E. R. Zunder, Sexual dimorphism in the dorsal root ganglia of neonatal mice identified by protein expression profiling with single-cell mass cytometry. Mol. Cell. Neurosci. 126, 103866 (2023).

56. S. Kumar, A. D. Kahle, A. B. Keeler, E. R. Zunder, C. D. Deppmann, Characterizing Microglial Signaling Dynamics During Inflammation Using Single-Cell Mass Cytometry. Glia. 73, 1022–1035 (2025).

57. J. Shi, W. Liu, A. Song, T. Sanni, A. Van Deusen, E. R. Zunder, C. D. Deppmann, Extrinsic apoptosis and necroptosis in telencephalic development: a single-cell mass cytometry study. Cell Death Differ. (2025), doi:10.1038/s41418-025-01594-5.

58. A. L. Van Deusen, S. Kumar, O. Y. Calhan, S. M. Goggin, J. Shi, C. M. Williams, A. B. Keeler, K. I. Fread, I. C. Gadani, C. D. Deppmann, E. R. Zunder, A single-cell mass cytometry-based atlas of the developing mouse brain. Nat. Neurosci. 28, 174–188 (2025).

59. K. I. Fread, W. D. Strickland, G. P. Nolan, E. R. Zunder, An updated debarcoding tool for mass cytometry with cell type-specific and cell sample-specific stringency adjustment. Pac. Symp. Biocomput. 22, 588–598 (2017).

60. E. R. Zunder, R. Finck, G. K. Behbehani, E.-A. D. Amir, S. Krishnaswamy, V. D. Gonzalez, C. G. Lorang, Z. Bjornson, M. H. Spitzer, B. Bodenmiller, W. J. Fantl, D. Pe’er, G. P. Nolan, Palladium-based mass tag cell barcoding with a doublet-filtering scheme and single-cell deconvolution algorithm. Nat. Protoc. 10, 316–333 (2015).

61. G. K. Behbehani, C. Thom, E. R. Zunder, R. Finck, B. Gaudilliere, G. K. Fragiadakis, W. J. Fantl, G. P. Nolan, Transient partial permeabilization with saponin enables cellular barcoding prior to surface marker staining. Cytometry A. 85, 1011–1019 (2014).

62. R. Finck, E. F. Simonds, A. Jager, S. Krishnaswamy, K. Sachs, W. Fantl, D. Pe’er, G. P. Nolan, S. C. Bendall, Normalization of mass cytometry data with bead standards. Cytometry A. 83, 483–494 (2013).

63. R. P. Schuyler, C. Jackson, J. E. Garcia-Perez, R. M. Baxter, S. Ogolla, R. Rochford, D. Ghosh, P. Rudra, E. W. Y. Hsieh, Minimizing batch effects in mass cytometry data. Front. Immunol. 10, 2367 (2019).

64. V. A. Traag, L. Waltman, N. J. van Eck, From Louvain to Leiden: guaranteeing well-connected communities. Sci. Rep. 9, 5233 (2019).

65. A. L. Van Deusen, S. M. Goggin, C. M. Williams, A. B. Keeler, K. I. Fread, I. Cheng, C. D. Deppmann, E. R. Zunder, A developmental atlas of the mouse brain by single-cell mass cytometry. BioRxiv (2022), doi:10.1101/2022.07.27.501794.

66. L. McInnes, J. Healy, N. Saul, L. Großberger, UMAP: uniform manifold approximation and projection. JOSS. 3, 861 (2018).

67. E. Becht, L. McInnes, J. Healy, C.-A. Dutertre, I. W. H. Kwok, L. G. Ng, F. Ginhoux, E. W. Newell, Dimensionality reduction for visualizing single-cell data using UMAP. Nat. Biotechnol. 37, 38–44 (2018).

